# A DNA-structured mathematical model of cell-cycle progression in cyclic hypoxia

**DOI:** 10.1101/2021.08.09.455605

**Authors:** Giulia L. Celora, Samuel B. Bader, Ester M. Hammond, Philip K. Maini, Joe Pitt-Francis, Helen M. Byrne

**Affiliations:** Mathematical Institute, University of Oxford, Oxford, UK; Department of Computer Science, University of Oxford, Oxford, UK; Oxford Institute for Radiation Oncology, Department of Oncology, University of Oxford, Oxford, UK

**Keywords:** cell-cycle, cyclic hypoxia, cancer, delay-differential equation, model calibration, model selection

## Abstract

New experimental data have shown how the periodic exposure of cells to low oxygen levels (*i.e.*, cyclic hypoxia) impacts their progress through the cell-cycle. Cyclic hypoxia has been detected in tumours and linked to poor prognosis and treatment failure. While fluctuating oxygen environments can be reproduced *in vitro*, the range of oxygen cycles that can be tested is limited. By contrast, mathematical models can be used to predict the response to a wide range of cyclic dynamics. Accordingly, in this paper we develop a mechanistic model of the cell-cycle that can be combined with *in vitro* experiments, to better understand the link between cyclic hypoxia and cell-cycle dysregulation. A distinguishing feature of our model is the inclusion of impaired DNA synthesis and cell-cycle arrest due to periodic exposure to severely low oxygen levels. Our model decomposes the cell population into four compartments and a time-dependent delay accounts for the variability in the duration of the S phase which increases in severe hypoxia due to reduced rates of DNA synthesis. We calibrate our model against experimental data and show that it recapitulates the observed cell-cycle dynamics. We use the calibrated model to investigate the response of cells to oxygen cycles not yet tested experimentally. When the re-oxygenation phase is sufficiently long, our model predicts that cyclic hypoxia simply slows cell proliferation since cells spend more time in the S phase. On the contrary, cycles with short periods of re-oxygenation are predicted to lead to inhibition of proliferation, with cells arresting from the cell-cycle when they exit the S phase. While model predictions on short time scales (about a day) are fairly accurate (i.e, confidence intervals are small), the predictions become more uncertain over longer periods. Hence, we use our model to inform experimental design that can lead to improved model parameter estimates and validate model predictions.

## 1. Introduction

The cell-cycle is one of the most fundamental and energy consuming processes in cell biology. It is divided into four phases: G1 (growth), S (DNA synthesis), G2 (growth and preparation for mitosis) and M (mitosis). Transitions between these phases are regulated by complex interactions between cellular pathways and external stimuli which normally act to maintain tissue homeostasis. These interactions are impaired in transformed cells, leading to uncontrolled proliferation, which is a key hallmark of cancer [33]. Cell-cycle dysregulation is further linked to the hallmarks of cancer because it promotes genetic instability, *i.e.*, increasing mutation frequency [45]. Loss of cell-cycle control also plays a significant role in the failure of standard treatments, such as chemotherapy and radiotherapy, where treatment relapse is driven by the emergence of small subpopulations of resistant cells. In normal cells, the DNA damage response (DDR) maintains genetic stability by promoting cell-cycle arrest to allow time for DNA repair or, when DNA damage is irreparable, by promoting cell death via the induction of apoptosis. The DDR is activated early during tumourigenesis as an anti-cancer barrier to oncogene activity and physiological stresses [7, 30]. However, continuous activation of the DDR results in selective pressure for the outgrowth of mutated cancer cells, with aberrant cell-cycle progression and apoptotic control [12, 30]. Exposure to insufficient oxygen levels, i.e., *hypoxia*, is a key driver of tumorigenesis. Hypoxic regions are commonly found in solid tumours [6, 29, 39, 40] as a result of uncontrolled cell proliferation and abnormal vascular structures. Exposure to severe levels of hypoxia (< 0.1% *O*_2_), which are only observed in pathophysiological conditions, leads to replication stress and consequent activation of DDR and the pro-apoptotic p53 tumour suppressor [27, 42, 47]. Further, conditions of less than 0.1% *O*_2_ are associated with resistance to radiotherapy and are therefore commonly referred to as radio-biological hypoxia (RH) [64].

Our study is motivated by evidence that the tumour microenvironment is characterised by highly dynamic oxygen levels. While chronic hypoxia affects tumour regions at a significant distance from vessels, acute/cycling hypoxia can occur close to, and far from, blood vessels, with periods ranging from seconds to hours/days [51]. While high frequency fluctuations are usually associated with vasomotor activity, processes that occur on longer time scales (*e.g.*, vascular remodelling) can generate cycles with longer periods [44]. Such periodic changes in the environment are known to cause inflammation, which promotes the survival of more aggressive forms of cancer that are resistant to standard treatments [4, 12, 15, 44, 52, 54].

It is possible to culture cells *in vitro* in controlled oxygen environments that partially mimic the fluctuating oxygen levels experienced by tumours *in vivo*. However, *in vitro* experiments are limited by the range of oxygen cycles that can be tested. By contrast, mathematical models can provide insight into a wider range of experimental conditions. Our aim in this work is therefore to develop a novel mechanistic model of the cell-cycle that can be combined with *in vitro* experiments, to increase our understanding of how cyclic hypoxia impacts the cell-cycle.

When exposed to radio-biological hypoxia (RH) *in vitro*, progress through S phase is inhibited due to a rapid reduction in the rate of DNA synthesis [27, 49]. This hypoxia-induced S phase block has been attributed to impaired functioning of the enzyme ribonucleotide reductase (RNR) [27, 46], which mediates *de novo* production of *deoxynucleotide triphosphates* (dNTPs). Since dNTPs are the building blocks of DNA, the decrease in dNTP levels in severe hypoxia causes DNA synthesis to stall. In contrast, milder levels of hypoxia (1-2% *O*_2_) do not impact dNTP levels [27]. Replication stress, which is defined as any condition impacting normal DNA replication, leads to activation of the DDR when local oxygen levels are sufficiently low. While cells can initially survive at such low oxygen levels, if severe conditions are prolonged then cell death occurs [49]. Alternatively, if oxygen levels are restored, cells can reenter the cell-cycle although they may accumulate DNA damage, likely associated with the accumulation of reactive-oxygen species (ROS) during reoxygenation [44]. Depending on the amount of damage sustained, activation of cell-cycle checkpoints causes cells to accumulate in the G2-phase and prevents damaged cells from entering mitosis [15, 31, 46]. Alternatively, if the reoxygenated cells are unable to repair the damage accumulated, they die. Periodic exposure to RH (cyclic hypoxia) may therefore lead to cell death and strong selective pressure for more aggressive clones with impaired checkpoint activation.

Common techniques for monitoring cell-cycle dynamics include flow cytometry and time-lapse microscopy using the fluorescent ubiquitination-based cell-cycle indicator (FUCCI). Both methods can be used to indirectly estimate how the fraction of cells in different stages of the cell-cycle changes over time, by measuring either DNA content (flow cytometry) or expression of cell-cycle related proteins (FUCCI). Since we have access to flow cytometry data, we build our mathematical model so that it can be calibrated against this type of data. As shown in the schematic in Fig. 1, flow cytometry can distinguish cells in different stages of the cycle sorting them according to their DNA content. While cells in the S phase are synthesising new DNA and, therefore, have a variable amount of DNA (from one to two copies), cells in the G1 and G2/M phases have exactly one and two copies of DNA, respectively. Although this method can fail to distinguish cells with similar DNA content (*i.e.*, cells in G1 and early S phases or, cells in the G2/M and late S phases), bromodeoxyuridine (BrdU) labelling can overcome this limitation. Since cells in the S phase incorporate BrdU into newly synthesised DNA, it is possible to distinguish these cells by measuring BrdU uptake [58]. The bivariate distribution of cells over DNA content and BrdU labelling can therefore be used to estimate cell-cycle distributions which are usually presented as time series data for the evolution of the fractions of cells in the G1, S or G2/M phases. These estimates can be further refined by measuring the expression of proteins involved with cell-cycle regulation, such as cyclin-dependent kinases [18, 58].

**Figure 1:**
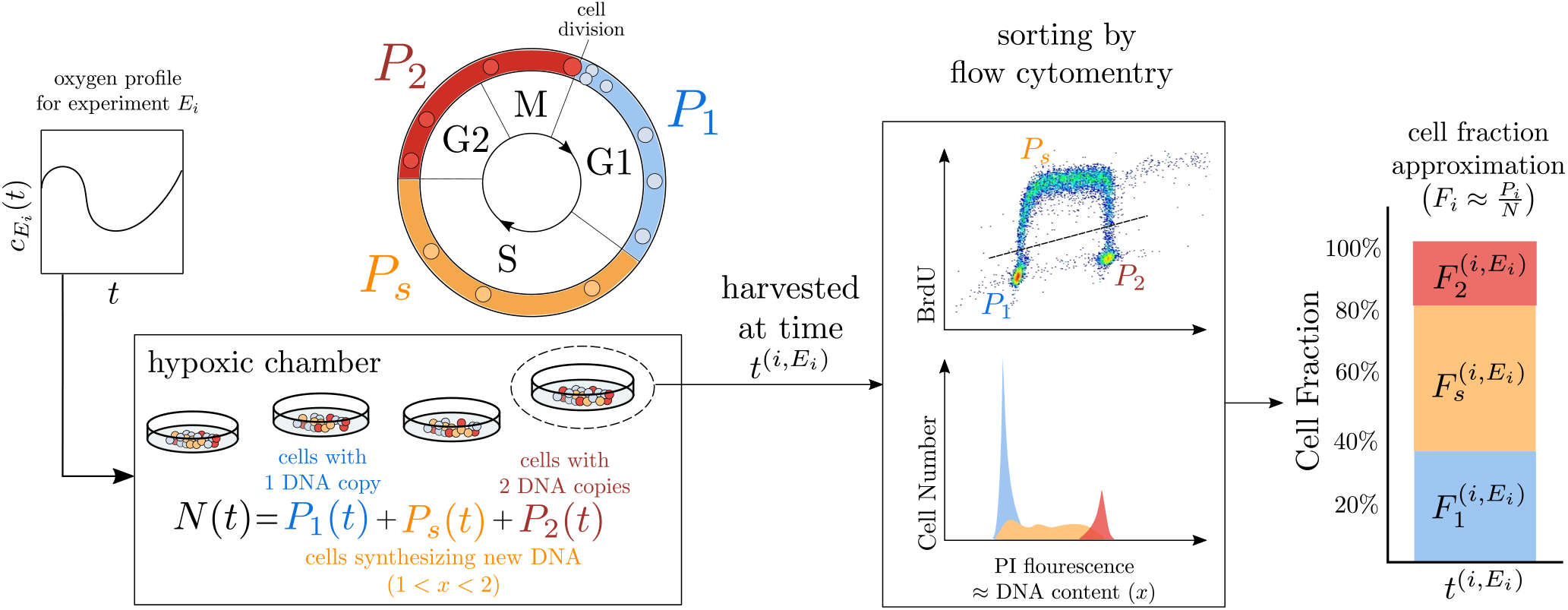
Schematic representation of flow cytometric data: at different time points 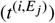 during experiment *E _j_* a subpopulation of *N* (the population of cells in the chamber) is harvested and sorted via flow cytometry according to their DNA content (PI fluorescence intensity), and BrdU uptake. This allows us to extract the fraction *F*_1_, *F_s_* and *F*_2_ of cells in the subpopulation with one, intermediate (i.e. in between one and two), and two copies of DNA, respectively. This is an approximation of the frequency of cells with different DNA content in the total population in the chamber *N*.

Several mathematical formalisms have been proposed to investigate cell-cycle evolution in normal ‘healthy’ conditions, as well as perturbed environments, for example in the context of drug development. These formalisms encompass discrete [26, 59], continuous (either deterministic [1, 8, 10, 25, 43, 54, 53] or stochastic [3, 63]) and hybrid approaches [2, 55]. When interested in the population scale, population balance (PB) models are often used [8, 10, 25, 43, 53, 56]. These take the form of age- and/or phase-structured models, where a structure variable is introduced to track progress through the cell-cycle. Several recent reports have focussed on developing compartment models of the cell-cycle, proliferation and migration which exploit time-resolved FUCCI data [16, 60, 61, 62]. When analysing flow cytometry data, instead, continuous structure variables are commonly used. For example, in a series of papers [8, 9, 10], Basse and coauthors developed cell-cycle models based on coupled partial differential equations in which cells are structured according to their DNA content. In these models, cells in the G1 and G2/M phases have constant DNA content (*x*), with *x* = 1 and *x* = 2, respectively, while *x* increases at a constant rate for cells in the S phase. Given a cell DNA content *x* at time *t*, and given its rate of DNA synthesis, it is possible to estimate the amount of time that a cell has spent in the S phase. However, information about the amount of time spent in the other phases of the cycle is lost. This shortcoming has motivated the development of age-structured models [11], in which the age of a cell in a certain phase of the cycle corresponds to the time it has spent in that phase. Models combining both structure variables (i.e., DNA and cell age) have also been proposed [17], but they are complex and difficult to validate against data. Further cellcycle specific properties such as size [25], or protein expression levels [18], can also be included in this framework. PB models can be extended to account for variability in compartment specific parameters, such as transition and/or death rates [25] and length of cell-cycle phases [8], to capture the effect of different drugs on cell-cycle progression. Several theoretical studies have investigated the effect of chronic exposure to hypoxia on cell-cycle arrest [1, 22] with particular emphasis on hypoxia-induced G1 arrest. At the population level, quiescence is usually represented by introducing an additional phase to the standard cell-cycle: the so-called ‘G0’ (or quiescent) phase [24, 32], which cells enter prior to committing to DNA synthesis (i.e, prior to entering the *S* -phase) when the local oxygen levels are too low (≈ 1% *O*_2_). While this extension can account for cell responses to chronic hypoxia, recent experimental findings [5, 27] suggest that it is not sufficient to account for cell behaviours in acute/cyclic hypoxia.

In this paper we develop a mathematical model to investigate how periodic (rather than constant) radio-biological hypoxia (RH) influences cell-cycle dynamics. In §2 we introduce the experimental data from [5] and summarise the biological mechanisms on which our model is based. In §3 we present our 5-compartment, DNA-structured, model which describes how the number of cells in each phase of the cycle evolves over time. Novel features of the model include a time and oxygen-dependent rate of DNA synthesis (here denoted by *v*(*t*)) and the introduction of two “checkpoint” compartments (*C*_1_ and *C*_2_), where cells arrest due to unfavourable environmental conditions or due to the accumulation of damage and stress. We further reduce our model to 4 ordinary differential equations coupled to a delay-differential equation, where the delay (τ_*S*_) represents the duration of the S phase and is not necessarily constant; rather it depends on the environment that the cells have encountered. In §4, we first consider model predictions in well-oxygenated conditions. Here we recover the well-known result of cells converging to a regime of balanced exponential growth. In §5, we explore model predictions in constant and cyclic RH, highlighting the distinct effects that these two types of hypoxia have on the cell-cycle dynamics. We first show that the model can replicate experimental data from [5] and then use it to investigate the cell-cycle dynamics for modes of oxygen fluctuations not yet tested experimentally. In §5.2, we introduce a class of models 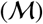 of decreasing complexity by neglecting some of the mechanisms (and complexity) included in the full model (such as arrest in *C*_1_ or *C*_2_). In §6 we compare these models using *Bayesian model selection*: we first calibrate our class of models against experimental data by using *Bayesian inference* methods and then identify the “best” model structure based on the *deviance information criteria*. This analysis reveals that the *C*_1_ and *C*_2_ checkpoint compartments are both necessary to describe the experimentally observed dynamics. In §7, we use the selected and calibrated model to predict cell responses to different oxygen environments when considering uncertainty in parameter estimates. We also explain how our results could inform the design of new experiments to validate the model and/or improve the accuracy of the parameter estimates. In §8 we conclude by summarising our results and outlining possible directions for future work.

## 2. Experimental motivation

This work is inspired by experimental data showing how the cell-cycle of RKO (colorectal cancer) cells changes when they are cultured *in vitro* as 2D monolayers and exposed to fluctuating oxygen levels [5]. As shown in Fig. 1, cells are cultured in chambers where the oxygen levels *c* = *c*(*t*) (where *t* is time) are carefully controlled and assumed to be spatially homogeneous. At prescribed time points, a subset of the cells is analysed using flow cytometry to estimate the fractions of cells in the G1, S and G2/M phases of the cycle, which we denote, respectively, by *F*_1_, *F_s_* and *F*_2_. Since each measurement requires cells to be harvested, measurement errors can be taken to be independent. In the absence of cell death, we have that by definition *F*_1_ + *F_s_* + *F*_2_ = 1, so only two of the three cell fractions are needed to fully characterise the cell-cycle dynamics.

As shown in Fig. 2, two experimental protocols are tested. At time *t* = 0, cells are exposed to either constant radio-biological hypoxia (*c*(*t*) ≡ *c_RH_* ≈ 0.1% *O*_2_, *t* > 0) or periodic cycles of radio-biological hypoxia (2 hr at *c* = *c_RH_* and 2 hr at 2% *O*_2_). Prior to both experiments, the cells are cultured in normoxia (21%*O*_2_) so that the measurements at time *t* = 0 contain information on this condition. In normoxia, and in absence of competition, the cells are typically in a regime of balanced exponential growth for which the cell fractions *F_m_* are stationary (i.e., they do not change over time). Hence a single set of measurements is sufficient to fully characterise the cell-cycle dynamics in these environmental conditions. We therefore divide the data into three different sets: *E*_0_ (normoxia), *E*_1_ (constant RH) and *E*_2_ (cyclic RH). The histograms in Fig. 2 summarise the data available from [5], obtained by averaging over between two and four replicates. As mentioned in §1, our focus is on fluctuating environments (i.e. the scenario *E*_2_), however data from *E*_0_ and *E*_1_ are also used to assist with parameter estimation.

**Figure 2:**
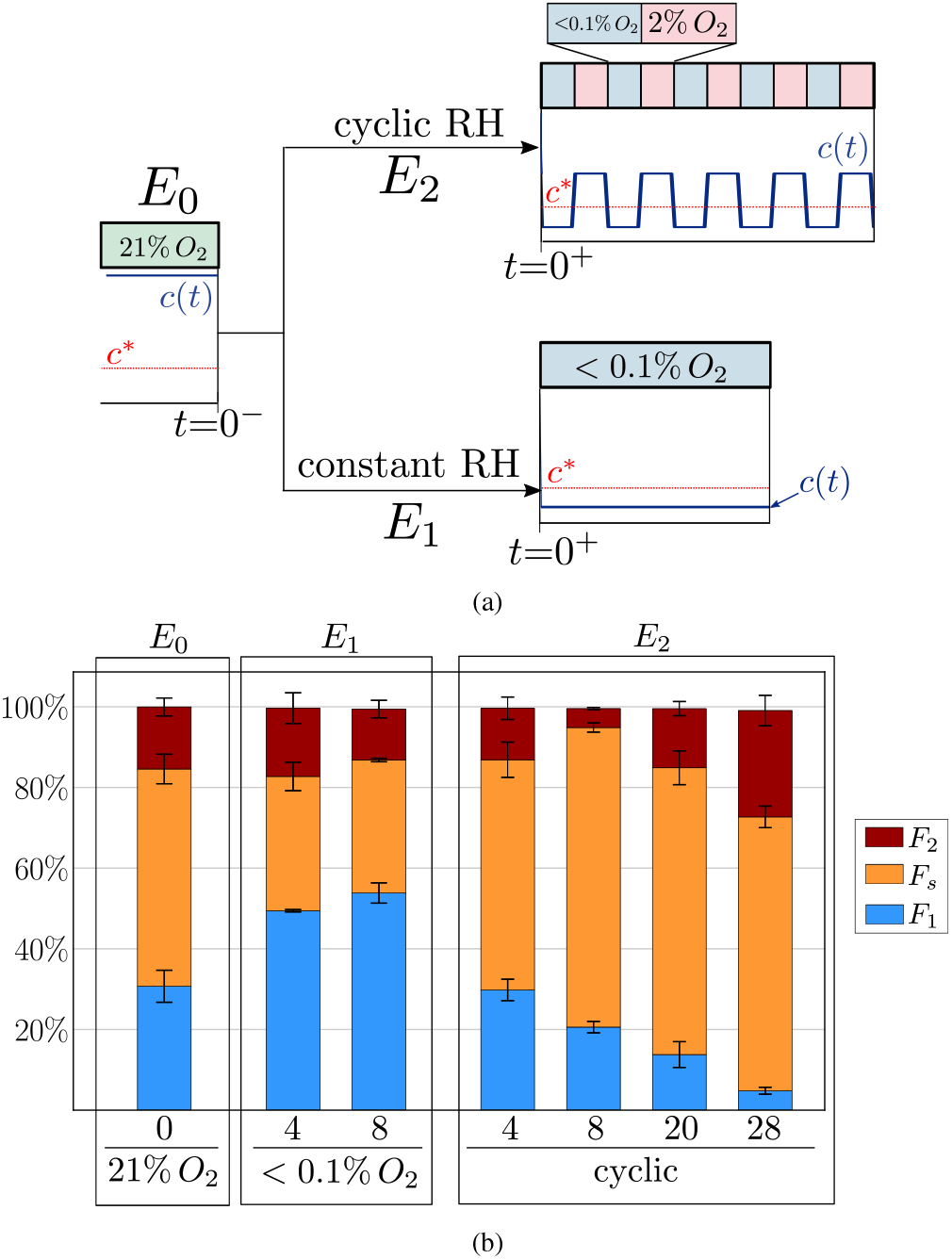
(a) Data are split into three sets corresponding to the three experiments : normoxia (*E*_0_), constant RH (*E*_1_) and cyclic RH (*E*_2_). We plot the evolution of the oxygen levels, *c*(*t*) (see blue curve), and compare it to the threshold *c*\* at which DNA synthesis is impaired (see red horizontal line). (b) Histogram summarising the data from [5]. At each time point we report the mean cell fractions estimated from multiple (between two and four) measurements. Error bars indicate one standard deviation in the estimated values. The complete data sets can be found in Appendix B.

The data in Fig. 2 show that both constant (*E*_1_) and cyclic RH (*E*_2_) redistribute the cells along the cycle. For experiment *E*_1_, cells tend to accumulate in the G1 phase (as seen by the increase in the fraction *F*_1_), whereas in experiment *E*_2_, we observe a decrease in the fraction *F*_1_. In this case, cells initially concentrate in the S phase (as seen by the increase in the fraction *F_s_*), while they accumulate in the G2/M phase at later times (see time *t* = 28 hr). For experiment *E*_1_ we only consider the early dynamics since on longer time scales (*t* > 20 hr) celldeath becomes significant. By contrast, cells can survive for longer periods in cyclic hypoxia, with cell death negligible for up to 28 hours.

As mentioned in the introduction, and based on current experimental evidence [4, 27, 49], three mechanisms may influence the cell-cycle dynamics under cyclic hypoxia: the reduction in the rate of DNA synthesis (Mechanism 1) and variation in the timing of the G1-S transition (Mechanism 2), both due to dNTP shortage and, the arrest of damaged cells in the G2 phase upon completion of the S phase (Mechanism 3). Here, we assume that there is an oxygen threshold, *c*\*, such that dNTP levels drop (due to the impaired activity of RNR enzyme) when oxygen levels fall below *c*\*. Based on the experiments in [27], we estimate that 0.1% *O*_2_ < *c*\* < 2% *O*_2_. We therefore fix *c*\* at an intermediate value (i.e., *c*\* ≈ 1%*O*_2_). Our aim is to develop a mathematical model that captures these three mechanisms and that can be used to investigate whether they can explain the experimental data in both constant RH (*E*_1_) and cyclic RH (*E*_2_) conditions.

## 3. The mathematical model

We propose a 5-compartment partial differential equation (PDE) model to describe cell-cycle dynamics in cyclic RH. For simplicity, we assume that the cells are in a well-mixed, spatially-homogeneous environment where the oxygen concentration *c* = *c*(*t*) is externally prescribed. We also assume that cell death is negligible since this is supported by the experimental observations.

As illustrated in Fig. 3, we subdivide the population into 5 compartments. Here *G*_1_ = *G*_1_(*t*) and *G*_2_ = *G*_2_(*t*) denote, respectively, the number of cells at time *t* in the G1 and G2/M phases of the cycle that are actively proceeding along their cycle. On the other hand, *C*_1_ = *C*_1_(*t*) and *C*_2_ = *C*_2_(*t*) are, respectively, the number of cells in the G1 and G2/M phases that are arrested at time *t*. We define the latter as checkpoint compartments and assume the rates of entry into and exit from these compartments change over time in response to the current and previous environmental conditions (i.e, oxygen levels). Finally, we structure cells in the S phase according to their DNA content, *x*, so that *S* = *S* (*x*, *t*) represents the number of cells with DNA content *x* at time *t*. Here *x* is a dimensionless variable corresponding to the relative DNA content of a cell, scaled so that 1 ≤ *x* ≤ 2. Cells start the S phase with *x* = 1 and exit it upon completion of DNA duplication with *x* = 2.

**Figure 3:**
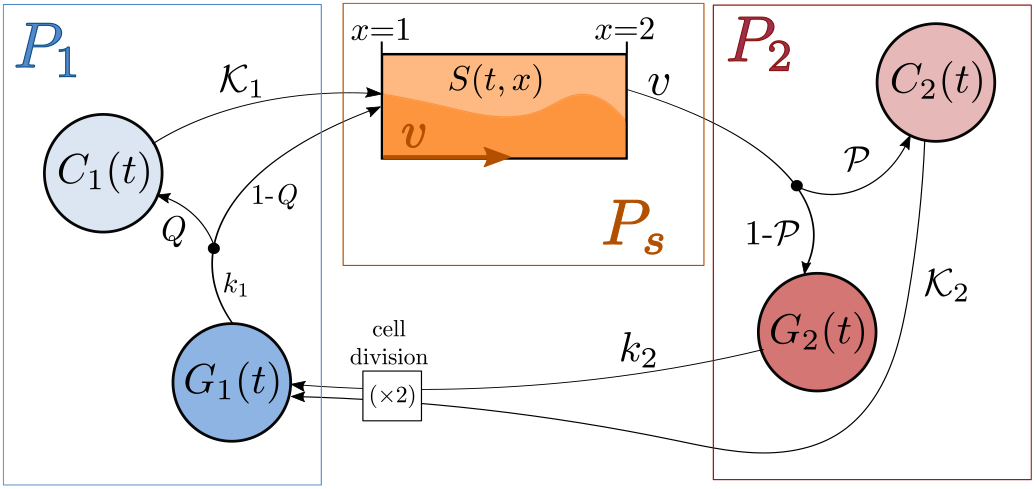
Schematic representation of the 5-compartment model of the cellcycle given by Equations (1). Here the two independent variables are time *t* and the cell DNA content *x* (measured in number of copies), while *P*_1_, *P_s_* and *P*_2_ are as defined in Fig. 1. The sub-population *P*_1_ groups cells in the *G*_1_ and *C*_1_ compartments, and similarly *P*_2_ comprises cells in the *G*_2_ and *C*_2_ compartments. Cells in the *G* compartments are progressing along the cell-cycle as usual, while the *C* compartments are “checkpoint” compartments where cells arrest. The black dots on the arrows correspond to redistribution of cell fluxes according to a given probability (for example in *P*_1_ the influx *k*_1_*G*_1_ is redistributed with probability *Q* into *C*_1_ and 1 − *Q* into *S*). To account for DNA synthesis during the *S* -phase we structure cells in the *S* -compartment according to their DNA content *x*, which is synthesised at velocity *v*.

Referring to the schematic in Fig. 3 and applying the principle of mass balance, we obtain the following system of equations for the time evolution of the model variables:

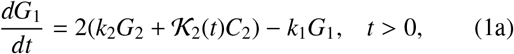

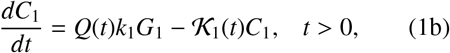

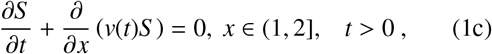

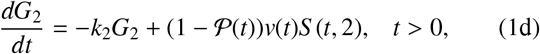

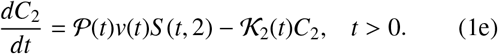

The factor of 2 in Eq. (1a) arises because cell division produces two daughter cells. The positive constants *k*_1_, *k*_2_ [hr^−1^] represent the rates at which cells leave *G*_1_, *G*_2_, respectively. In Eq. (1c), we account for DNA synthesis by assuming that cells in the S compartment are advected along the *x*-axis at velocity *v*(*t*) > 0 [hr^−1^] (i.e., they produce DNA at a rate *v*(*t*)). In Eq. (1b), we assume that cells exiting the *G*_1_ compartment arrest (*i.e.* enter the *C*_1_ compartment) with probability *Q* ∈ [0, 1], while they proceed to enter the S phase with probability 1 − *Q*. We further assume that arrested cells re-enter the cycle at rate 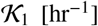. In a similar manner, in Eq. (1e) we assume that cells exiting the *S* compartment arrest in *C*_2_ with probability 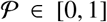 and arrested cells may re-enter the cycle at a rate 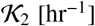. We account for the effect that different oxygen levels have on cell behaviour by allowing certain model param-eters (specifically 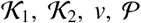 and *Q*) to vary over time in response to the oxygen levels *c*. The boundary condition for Equation (1c) is derived by applying conservation of cell flux at *x* = 1. Under the assumption that *v*(*t*) > 0 for all *t* > 0, we find:

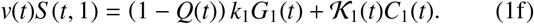

We close Eqs. (1a)–(1f) by imposing the following initial conditions:

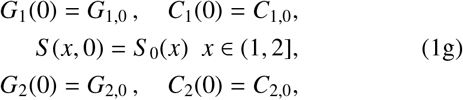

where *G_i_*_,0_, *C_i_*_,0_ are constants and *S* _0_(*x*) is a prescribed function. Since cell numbers can not be negative, we have that *G_i_*_,0_ ≥ 0, *C_i_*_,0_ ≥ 0 and *S* _0_(*x*) ≥ 0 for all *x* ∈ [1, 2].

In order to compare the model output to flow cytometry data, we need to express cell fractions (see Fig. 1) in terms of our model variables. As shown in Fig. 1, cells can be divided in three sub-populations depending on the cell-cycle phase they are in: *P*_1_, *P_s_* and *P*_2_. Cells in G1 belong to *P*_1_, cells in G2/M to *P*_2_ and cell in S to *P_s_*. The total number of cells, *N* = *N*(*t*), is then obtained by summing the number of cells in each subpopulation:

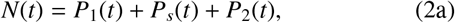

where (as illustrated in Fig. 3)

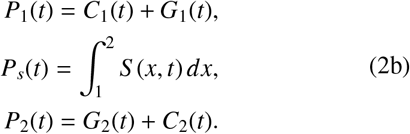

Differentiating Eq. (2a) with respect to time and using (1), we find that *N*(*t*) satisfies:

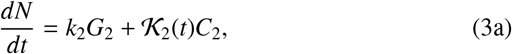

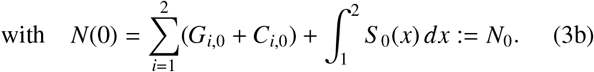

The population proliferation rate, *ω*, is given by:

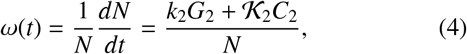

while the cell fractions *f_m_* are defined by:

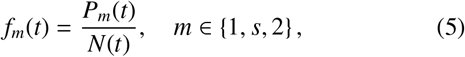

and correspond to the probability that a cell randomly chosen from the total population belongs to the sub-population *P_m_*. We note hat the fractions *f_m_* are not independent; Eq. (2) implies that 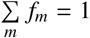. We further introduce the distribution *s* = *s*(*x*, *t*):

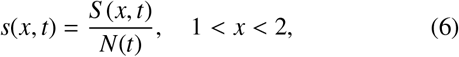

which corresponds to the probability that a cell in the subpopulation *P_s_* has a DNA content *x* ∈ [1, 2] at time *t*. A summary of the model variables is given in Table 1.

A key feature of our model is that the velocity *v* is assumed to depend on the oxygen levels *c* in order to account for impaired DNA synthesis at low oxygen levels (*c* < *c*\*). In particular, we propose the following piece-wise linear ODE to describe how the advection velocity *v* adapts to changes in local oxygen levels *c*(*t*):

**Table 1:** Summary of the model variables in Eqs. (1) and (8).

|  | Description |
| --- | --- |
| $t$ | time [hr] |
| $x$ | copies of DNA in a cell ($1.0 \leq x \leq 2.0$) |
| $c$ | externally controlled oxygen levels [%] |
| $G_1(t)$ | number of cells in $P_1$ active in the cycle |
| $C_1(t)$ | number of cells in $P_1$ that are arrested |
| $S(x, t)$ | number density of cells in $P_s$ with DNA content $x$ |
| $G_2(t)$ | number of cells in $P_2$ active in the cycle |
| $C_2(t)$ | number of cells in $P_2$ that are arrested |
| $N(t)$ | total cell number |
| $f_m(t)$ | predicted fraction of cells ($f_m \in [0, 1]$) in the $P_m$ sub-population $m \in \{1, s, 2\}$ |
| $F_m(t)$ | experimentally measured fraction of cells ($F_m \in [0, 1]$) in the $P_m$ sub-population $m \in \{1, s, 2\}$ |
| $s(x, t)$ | DNA distribution of cells in the $P_s$ sub-population |
| $\tau_S(t)$ | duration of the S phase for cells that exit the S compartment at time $t$ [hr] |
| $\omega(t)$ | population proliferation rate [$\text{hr}^{-1}$] |

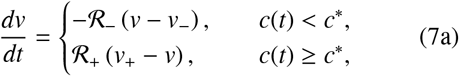

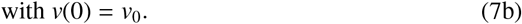

In Eqs. (7), 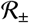, *v*_0_ and *v*_±_ are positive constants, while *c*\* is the threshold oxygen level below which RNR activity is impaired (see §2 for details). Here *v*_+_ and *v*_−_ represent, respectively, the equilibrium velocities in well-oxygenated (*c* > *c*\*) and severely hypoxic (*c* < *c*\*) environments and 0 < *v*_−_ < *v*_+_. It is straight-forward to show that if *v*_0_ ∈ [*v*_−_, *v*_+_], then *v*(*t*) ∈ [*v*_−_, *v*_+_] at all times *t* > 0, so that *v*_+_ and *v*_−_ can also be viewed, respectively, as the maximum and minimum rates of DNA synthesis. When oxygen levels drop below the critical value *c*\*, the rate of DNA synthesis decreases (i.e. *dv*/*dt* 0 for *c* < *c*\*), capturing the effect of dNTP shortage. As observed in [27], cells maintain a minimum level of DNA synthesis even after prolonged exposure to severe hypoxia (*c* < *c*\*); we therefore assume that *v*_−_ > 0, which guarantees that *S* (*t*, 1), as defined by Eq. (1f), remains finite. Once *c* > *c*\*, dNTP levels are restored and the rate of DNA synthesis increases (i.e., *dv*/*dt* > 0), towards its maximum value *v*_+_. The rates 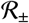 are assumed constant for simplicity although, in practice, they may odxepyegnedn olenvels. In the absence of suitable data to identify these dependencies, we proceed with the simplest form that captures the currently available data and postpone investigation of more complex functional forms to future work.

We introduce the checkpoint compartment *C*_1_ to account for transient arrest of cells in the *P*_1_ phase due to dNTP shortage. The rates of transition into (*Q*) and out of 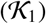 the compartment depend on current oxygen levels. For simplicity, we adopt the following functional forms:

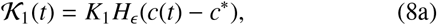

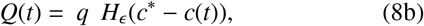

where *K*_1_ and *q* are positive constants with *q* ∈ [0, 1] and *c*\* is the oxygen threshold introduced above. The function *H*_*ϵ*_ is the standard continuous approximation of the Heaviside function:

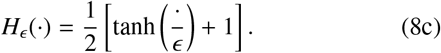

where 0 < *ϵ* ≪ 1 is a small parameter, here set to *ϵ* = 0.01% oxygen. Note that Eqs. (8a)–(8b) ensure that cells arrest when *c* < *c*\* and that they resume cycling by entering the S phase once *c* > *c*\*. We considered alternative functional forms for 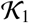 and *Q* (such as assuming them to be proportional to *v*) but found that the resulting model was unable to capture the experimentally observed dynamics (results not shown).

In Eq. (7), prolonged or frequent exposure to RH slows DNA replication and, therefore, increases the time τ_*S*_ that cells spend in the S phase. More precisely, we denote by τ_*S*_ = τ_*S*_ (*t*) the amount of time a cell exiting the S phase at time *t* has taken to complete DNA synthesis (we will explain how τ_*S*_ (*t*) is computed in §3.1). Consequently, the larger τ_*S*_ , the more cells have been damaged during the S phase due to re-oxygenation and replication stress (in RH). Since the accumulation of damage regulates the arrest of cells in the G2 phase (here captured by cells transitioning into the *C*_2_ compartment), we assume that the probability, 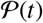, of arrest in G2 increases with τ_*S*_ (*t*). In particular, we assume that, when cells exit the S phase, they arrest with a probability 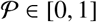 where:

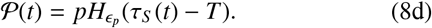

In Eq. (8d), we fix the small parameter *ϵ*_*p*_ so that *ϵ*_*p*_ = 0.1 hr, while *p* ∈ [0, 1] and *T* > 0 are unknown parameters. Here *p* is the maximum probability that a cell enters the *C*_2_ compartment while *T* captures the critical duration of the *S* phase after which cells are likely to arrest in *C*_2_. Based on Eq. (7), the timescale for completing the *S* phase, τ_*S*_, satisfies 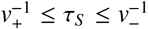. When oxygen levels are sufficiently high, (i.e. *c*(*t*) > *c*\* for all *t*), neglecting an initial transient in the case *v*_0_ ≠ *v*_+_, we have that 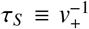. Since we do not expect cells to arrest in an oxygenrich environment we require 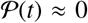 when 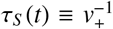. Consequently we set 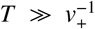. As for the *C*_1_ compartment, we assume that, once oxygen levels rise above the critical value *c*\*, cells can repair any damage they have accumulated and re-enter the cell-cycle at a rate 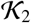:

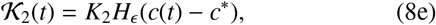

where *K*_2_ is a non-negative constant and *ϵ* = 0.01% oxygen as in Eq. (8a).

### 3.1. Model reduction to a system of Delay Differential Equations (DDEs)

Here we show how Eqs. (1) can be rewritten as a system of ODEs coupled to a (state-dependent) DDE with a non-constant delay τ_*S*_ (*t*) which we view as a state variable. Given that the velocity *v* is always positive, shocks can not form and a straight-forward application of the *method of characteristics* to Eq. (1c) yields

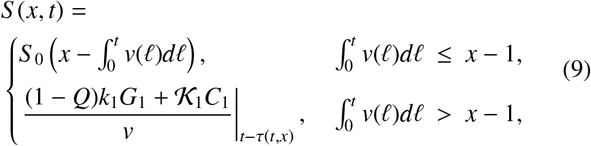

where the function τ = τ(*t*, *x*) is implicitly defined by:

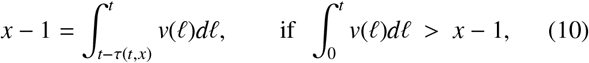

and indicates the amount of time a cell with DNA content *x* at time *t* has spent in the S compartment.

As illustrated in Fig. 4, the condition 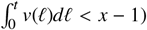 divides the (*t*, *x*)-plane into two regions depending on whether the characteristics propagate from the boundary data curve (orange curve) or the initial data curve (green curve). When *t* > *t*\* (i.e. when *S* (2, *t*) is influenced by the boundary data) the total time spent in the *S* compartment is given by τ_*S*_ (*t*) = τ(*t*, 2). Using Eq. (10), we find that τ_*S*_ is implicitly defined by:

**Figure 4:**
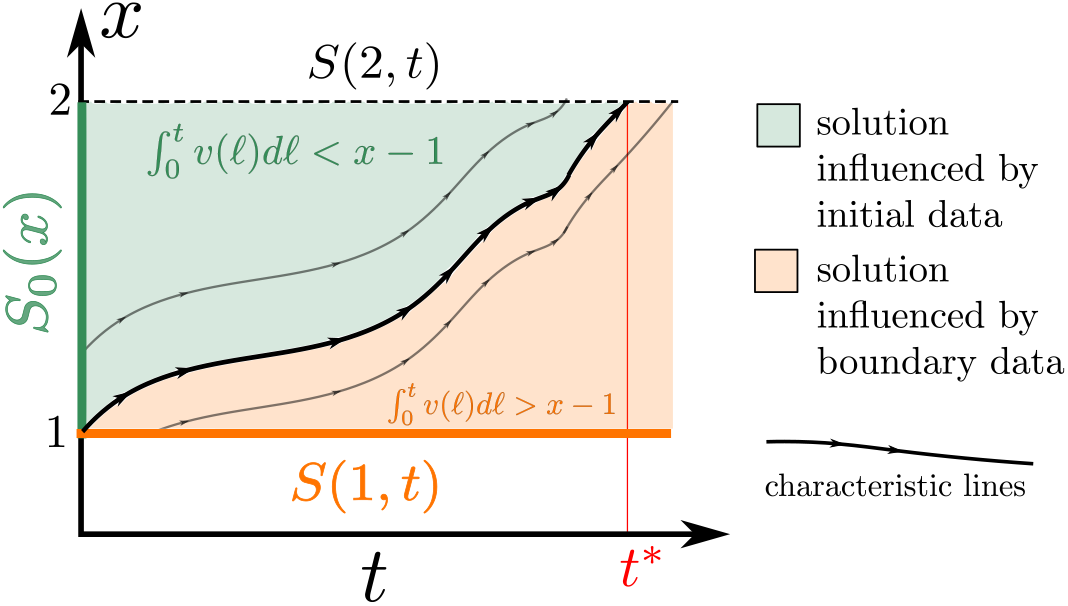
Schematic illustration of the characteristic curves in the (*t*, *x*) plane. The bold curve divides the plane into two distinct regions: in the green region (where 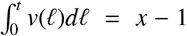) the solution *S* (*x*, *t*) is determined by the initial data; in the orange region (where 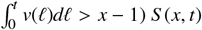) is instead determined by the boundary data.

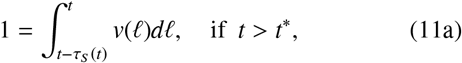

where *t*\* is defined implicitly by the integral equation

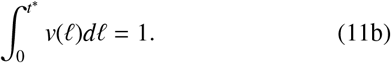

However, more information about the behaviour of cells prior to the beginning of the experiment is needed to estimate τ_*S*_ (*t*) for 0 *t* < *t*\*. Let us consider a cell that exits the *S* phase at time *t* < *t*\*. Then, at time *t* = 0, its DNA content was:

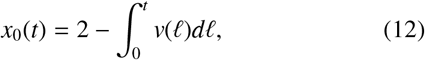

as we can write τ_*S*_ = *t* + τ_0_, where *t* denotes the time spent in the *S* compartment since the beginning of the experiment and τ_0_ the time spent in the S phase prior to the beginning of the experiment. The functional form of τ_0_ will depend on the conditions in which the cells were grown for *t* < 0. It is therefore part of the initial data that must be specified in order to fully define the model. In §4, we will show that if cells are cultured in an oxygen-rich environment for *t* < 0, then τ_0_ depends only on the DNA content *x*_0_(*t*) (*i.e.* τ_0_(*t*) = τ_0_(*x*_0_(*t*))). In summary we have:

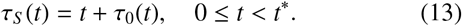

Differentiating Eq. (11a) with respect to time, and given Eq. (13), we deduce that the variable τ_*S*_ (*t*) satisfies:

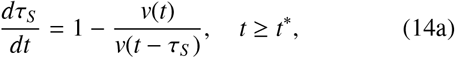

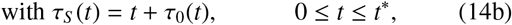

### 3.2. Summary of the cell-cycle model with delay

Before presenting the full model, we perform the following re-scaling:

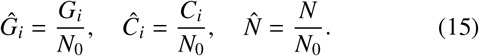

Since our model is linear, this re-scaling does not affect the form of the governing equations; it only alters the initial conditions. Consequently we can determine the system dynamics without data on the initial cell number. This is helpful since we only have experimental data on the initial cell fractions, while *N*_0_ is unknown. We further assume that cells are initially actively cycling (*i.e.* none of them is arrested) and DNA synthesis is proceeding at maximum speed so that:

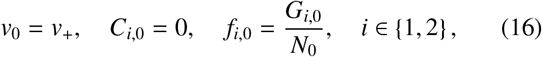

where *f_i_*_,0_, with *i* ∈ {1, 2}, denote the cell fractions at time *t* = 0. As discussed in §4, this is a reasonable assumption if cells are cultured in oxygen-rich environments prior to the start of the experiments.

Applying Eq. (15) to Eqs. (1)–(3), using Eqs. (7)–(8), (1c) and (14) and dropping the hat notation, the governing equations become

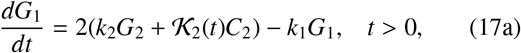

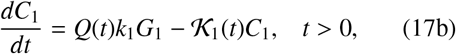

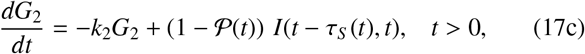

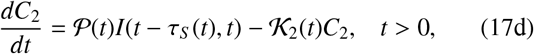

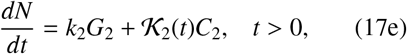

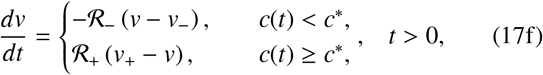

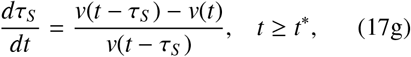

wherein

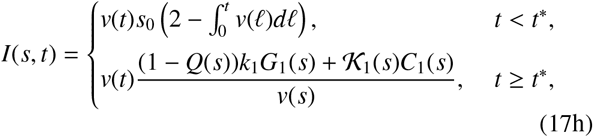

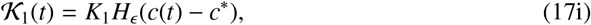

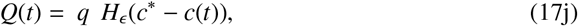

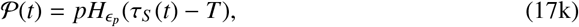

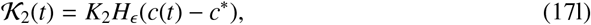

and

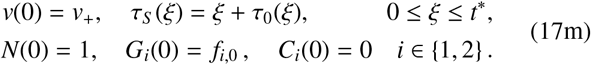

In Eqs. (17h)–(17m) it remains to specify the distribution *s*_0_ = *S* _0_/*N*_0_ and the function τ_0_ (see §3.1). The former is subject to the constraint:

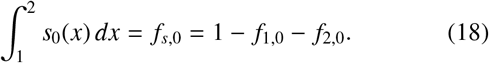

In Fig. 5, we present a schematic summary of the model (as in Fig. 3) where the *S* -compartment is replaced by the time delay τ_*S*_.

**Figure 5:**
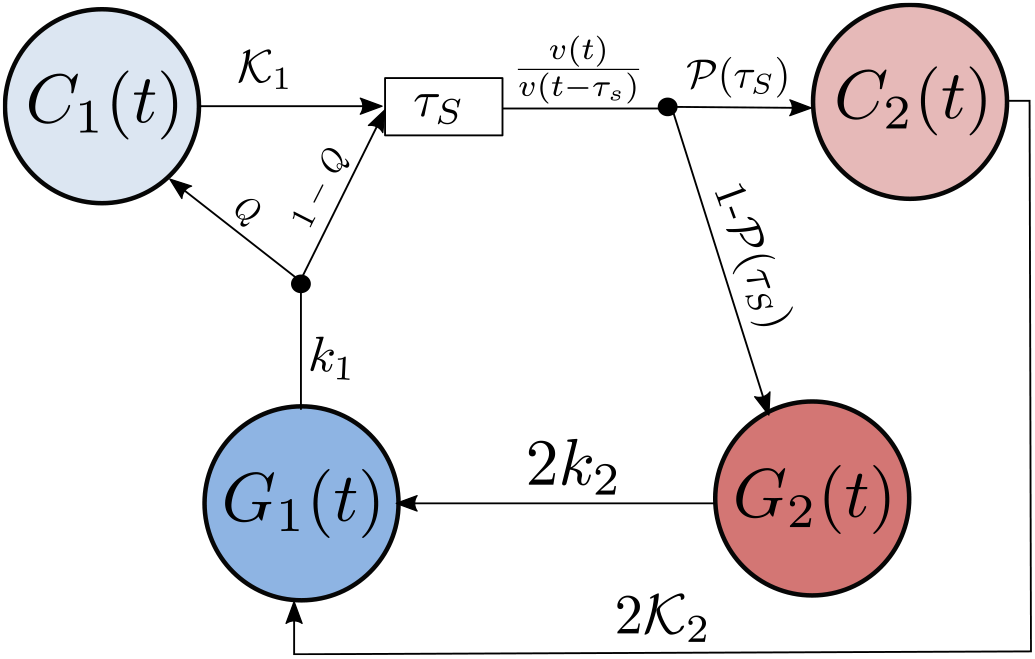
Schematic representation of the 4-compartment model with delay (see equations (17)) derived by solving Eq. (1) via the method of characteristics. The black nodes correspond to the conservation of the cell fluxes, where the input is redistributed with probabilities *P*(τ_*S*_) (*Q*(*c*)) and 1 − *P*(τ_*S*_) (1 − *Q*(*c*)) in *C*_2_ (*C*_1_) and *G*_2_ (*G*_1_), respectively.

As listed in Table 2, our model has 12 parameters. As discussed in §2, we fix *c*\* = 1% oxygen in line with experimental evidence. Further, based on the experimental estimates of the rate of DNA synthesis in [49], we expect *v*_−_ to be small. Since the limit *v*_−_ → 0 is non-singular, it can be shown that the solution is not sensitive to the precise value of *v*_−_ provided that *v*_−_ ∼ *O*(10^−3^) (results not shown). We therefore fix *v*_−_ = 0.005 hr−1, which is sufficiently small to exhibit the correct qualitative behaviour. With *c*\* = 1 and *v*_−_ = 0.005, there are ten unknown parameters, which we split into two classes. While **θ**=[*k*_1_, *v*_+_, *k*_2_] comprises parameters associated with oxygen-independent mechanisms, 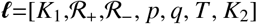 contains parameters associated with oxygen-dependent mechanisms. Given the large number of unknown parameters and the expected variability for different cell lines, we here focus on the RKO cancer cell line and the data in Fig. 2 to estimate the model parameters. Discussion on how we estimate the model parameters is postponed to §6. First, we present some characteristic predictions of our model obtained by solving Eqs. (17) numerically using the Python *ddeint* package to integrate delay-differential equations. As we will justify in §5.2, we assume the *C*_2_ checkpoint to be irreversible by setting *K*_2_ = 0. This is likely to be the case for the RKO cancer cell line considered, but may not be true for other cell lines, particularly if they are p53-deficient, given the role of p53 in mediating G2 arrest in several cancer cells [57].

**Table 2:**
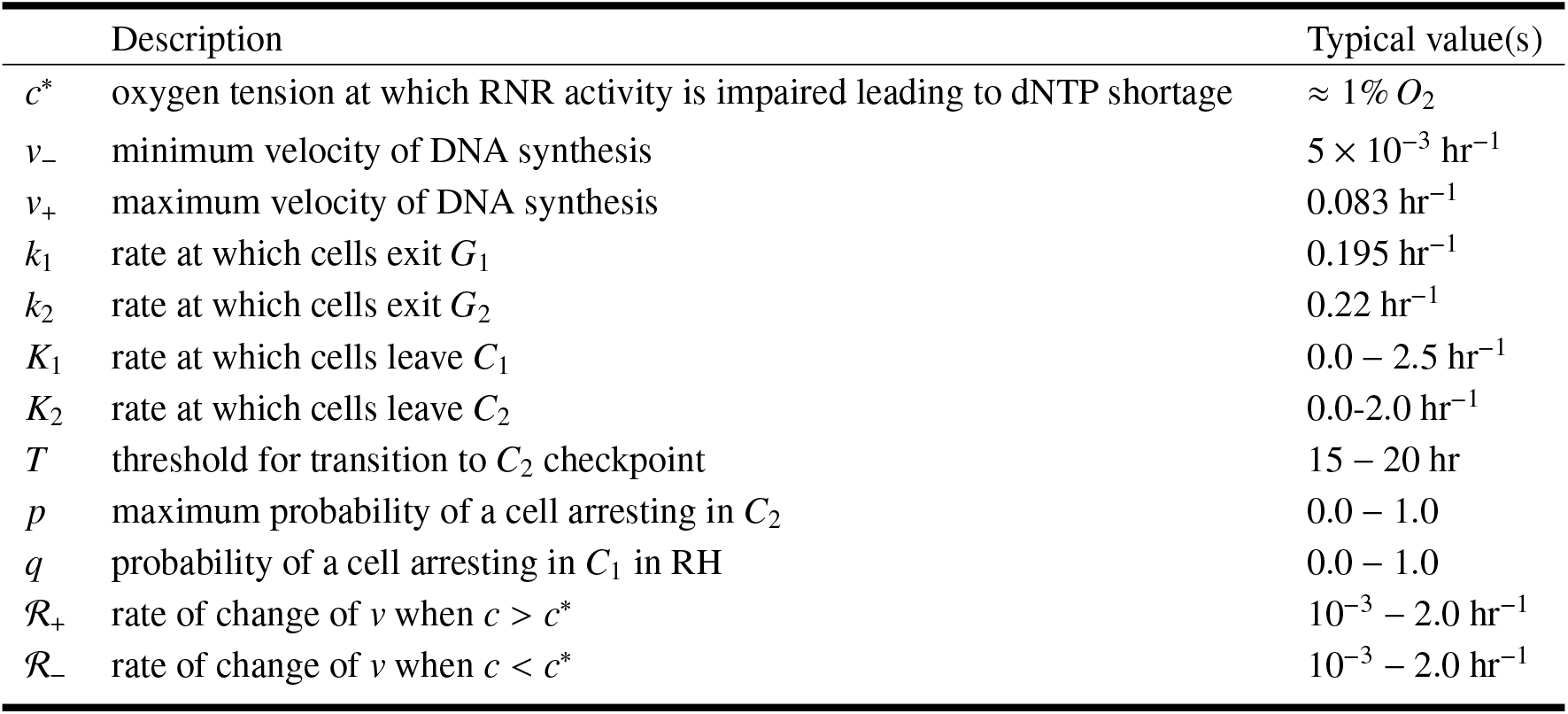
Summary of the parameters that appear in Eqs. (1)–(8), together with their typical values for the RKO cancer cell line.

## 4. Cell-cycle progression in a static normoxic environment

Let us first consider the case in which cells are exposed to a constant, oxygen-rich environment (i.e., 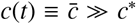). Given Eqs. (17f)–(17g) and (17m), we have that *v* ≡ *v*_+_ and 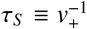. In this case, *Q* ≡ 0, *P* ≡ 0 and *C*_1_(*t*) = *C*_2_(*t*) ≡ 0 for all *t* > 0, so that there are no arrested cells. We conclude that when 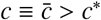, Eqs. (17) reduce to the following system:

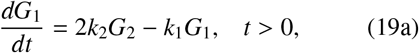

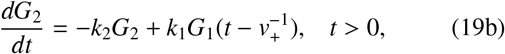

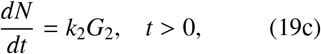

subject to

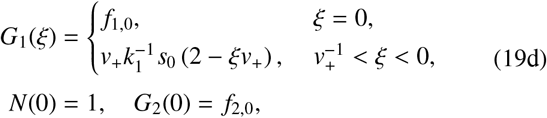

where *s*_0_ satisfies (18). We note that Eqs. (19) i s analogous to models previously proposed in the literature, such as the model by Basse et al. [8, 9] when dispersion is neglected. Since Eqs. (19) are linear with a constant delay, they can be solved exactly via superposition of exponential functions 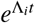 where Λ_*i*_ (*i* = 1, 2, …) are the complex roots of the characteristic polynomial (see Eq. (A.3) in Appendix A). In the case of DDEs, the characteristic polynomial is a transcendental equation with an infinite number of roots so that the computation of Λ_*i*_ is non-trivial. To investigate the transient dynamics, it is therefore more convenient to solve Eqs. (19) numerically. In Fig. 6 we present numerical solutions for two sets of initial conditions: cells are initially synchronised in either the *G*_1_ (Fig. 6(a)) or *G*_2_ (Fig. 6(b)) compartment. This corresponds to setting *s*_0_ ≡ 0 with *f*_1,0_ = 1 and *f*_2,0_ = 0 (for panel (a)), or *f*_2,0_ = 1 and *f*_1,0_ = 0 (for panel (b)).

**Figure 6:**
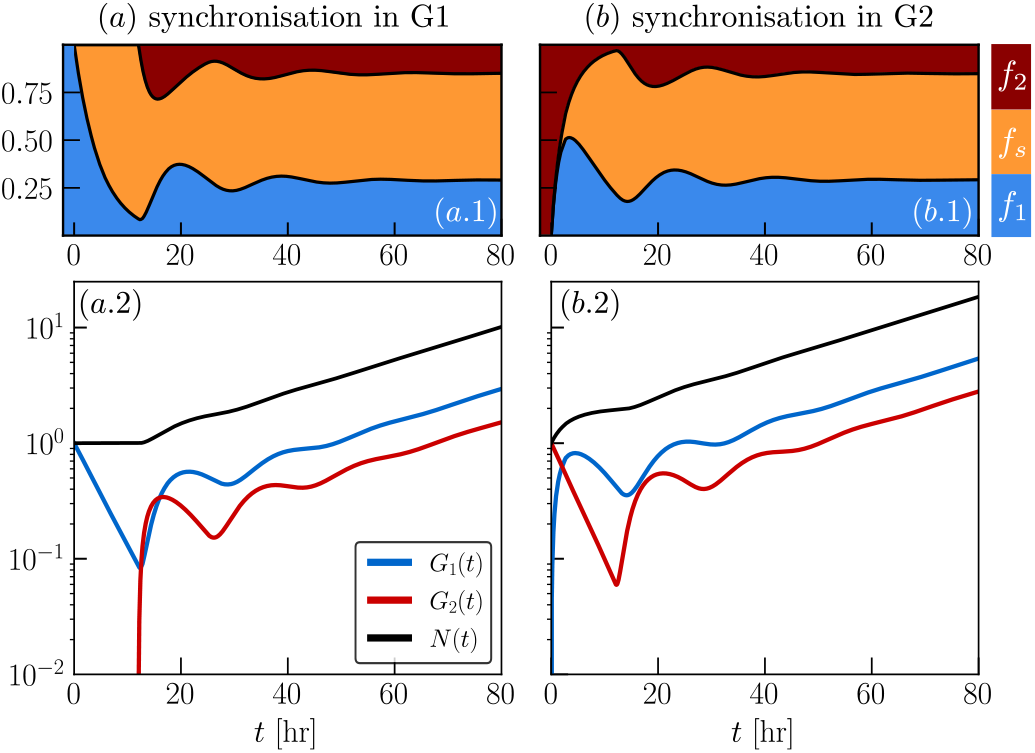
Numerical simulations of Eqs. (19) for two sets of initial conditions: (a) *s*_0_ ≡ 0, *f*_1,0_ ≡ 0 and *f*_2,0_ = 1; (b) *s*_0_ ≡ 0, *f*_1,0_ ≡ 1 and *f*_2,0_ = 0. For both simulations, the parameters *k*_1_, *k*_2_ and *v*_+_ are set as in Table 4. In panels (a.1) and (b.1), we plot the cell-cycle dynamics with cumulative plots of the cell fractions *f_m_* with *m* 1, *s*, 2; in panels (a.2) and (b.2) we show instead the predicted time evolution of the model variables *G*_1_, *G*_2_ and *N* on a semilogarithmic scale.

As shown in Fig. 6, the evolution of the cell fractions *f_m_* in panels (a.1) and (b.1) differs only up to time *t* ≈ 20 hr; after this first transient the cell fractions evolve to constant values, denoted by 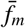, which are independent of the initial conditions. When looking at Fig. 6(a.2)-(b.2), we see that the evolution of *G*_1_, *G*_2_ and *N*, differs even at long times. From time *t* > 20 hr, the variables have a similar qualitative behaviour, but their values remain higher for scenario (b) than for scenario (a). While cells in scenario (b) start proliferating at the beginning of the simulations, cells in scenario (a) are delayed since they need to complete the *S* -phase before they can replicate. Once cells enter the *G*_2_ compartment, we see an increase in the cell number *N*. We also note that from time *t* > 60 hr, for both scenarios, *N*(*t*), *G*_1_(*t*) and *G*_2_(*t*) increase exponentially at a constant rate. This agrees with the results in Fig. 7, which show that the population proliferation rate *ω*(*t*) (see Eq. (4)) asymptotes to a constant value λ for both sets of simulations.

**Figure 7:**
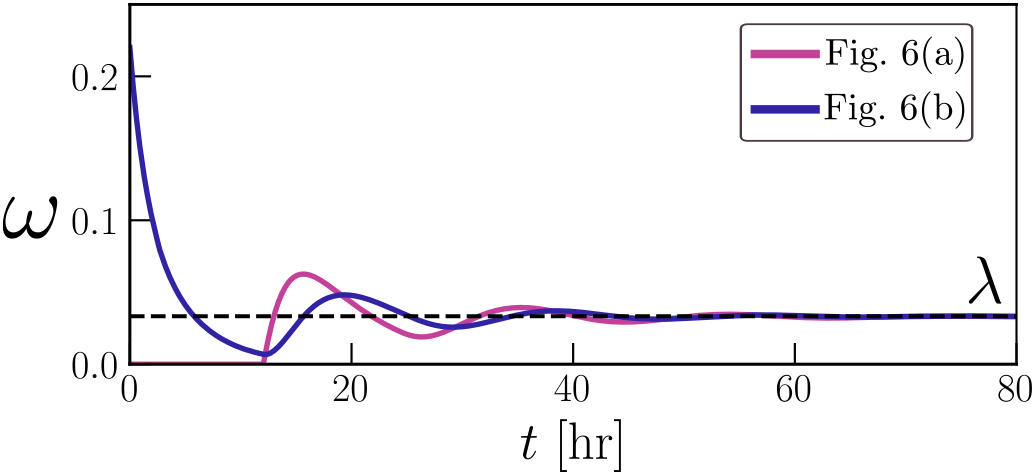
Comparison of the population proliferation rate *ω* (see Eq. (4)) for the two scenarios in Fig. 6. We see that on the long time scale, the proliferation rate settles to a constant value, λ, independently of the initial conditions chosen.

Fig. 8 shows the evolution of the distribution *s*(*x*, *t*). We note that in Fig. 8(a), cells are initially highly synchronised in the S compartment, with the formation of a front that propagates at velocity *v_max_* (note the steep gradient in the profiles at time *t* = 5 and *t* = 10). This is because of the discontinuity in the initial data for *G*_1_ (when *f*_1,0_ = 1 and *S* _0_ ≡ 0, *G*_1_(*ξ*) in Eqs. (19) is discontinuous at *ξ* = 0). The discontinuity propagates along the *x* axis but it quickly smooths out, due to the de-synchronisation of cells in the *G*_1_ and *G*_2_ compartments. By contrast, in Fig. 8(b), there is no discontinuity in the initial data for *G*_1_ and therefore the profile of *s* is smoother. Despite these large differences in the distributions at early times, *s* eventually evolves to the same stationary distribution (in Fig. 8, the curves for *t* = 60 and *t* = 80 are almost indistinguishable) and the time scales required to approach the stationary distribution for the two initial conditions are comparable.

**Figure 8:**
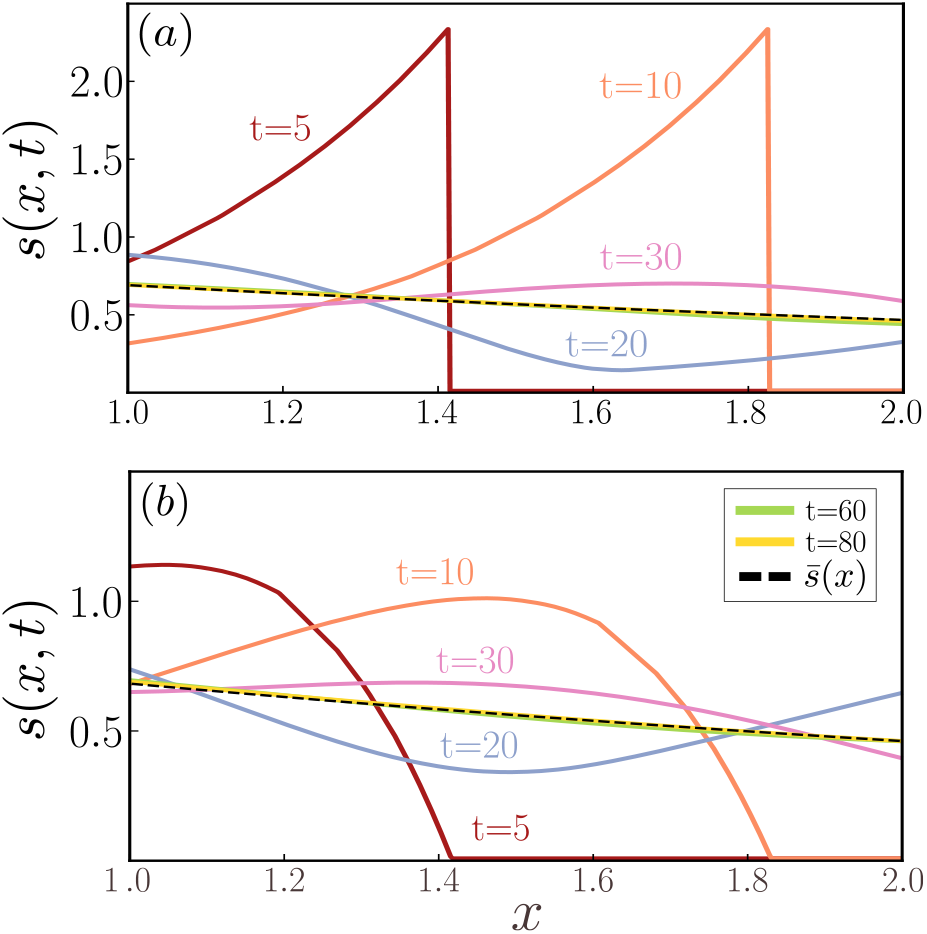
Evolution of the distribution of cells in *P_s_* with respect to *x*. We estimate the distribution *s*(*x*, *t*) by numerically solving Eq. (9) for the two cases in Fig. 6: (a) initial synchronisation in *G*_2_ (as in Fig. 6a) and (b) initial synchronisation in *G*_1_ (as in Fig. 6b). We compare the long time behaviour with the analytically computed solution from the phase stationary solution (see black dashed line). Here the colour scheme in (a) and (b) is the same.

Following the notation introduced in [54], we term the asymptotic solution of the model a *phase stationary solution* (PSS) to indicate that, in this regime, the cell fractions *f*_1_, *f_s_* and *f*_2_, and the distribution *s*(*x*, *t*) remain constant in time. This is similar to predictions from other models in the literature [8, 10, 13, 21, 54] in the context of unperturbed growth; this regime is usually referred to as balanced, or asynchronous, exponential growth [9, 14]. As mentioned previously, we can write the solution to Eqs. (19) as a superposition of exponential functions 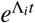. At long times, the behaviour is dominated by the exponential whose eigenvalue Λ_*i*_ has the largest real part (here denoted by λ). It is possible to prove that this eigenvalue λ is real and positive (see Appendix A for details). We conclude that for *t* ≫ 1 the system approaches a regime of exponential growth (as observed in the numerical results in Fig. 6) in which the model variables take the form:

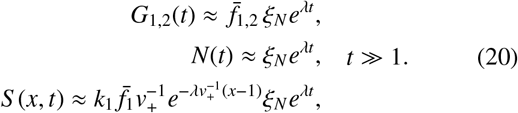

where 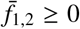 and *ξ*_*N*_ > 0 are constant. Substituting Eqs. (20) into Eqs. (5), we obtain that for *t* ≫ 1, 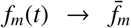 for *m* ∈ {1, *s*, 2}. Therefore 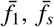 and 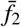 indicate, respectively, the stationary values of cell fraction in the G1, S and G2/M phase of the cycle. Focusing on the distribution of cells *s*(*x*, *t*), this converges to the stationary distribution 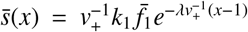, which is monotonically decreasing in *x* (see black line in Fig. 8). This indicates that, for *t* ≫ 1, cells in S phase are more likely to be starting DNA synthesis (with *x* ≈ 1) rather than close to completing it (with *x* ≈ 2). We note that the longer the duration of the S phase, or equivalently the smaller *v*_+_, the steeper is the 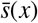 curve and, therefore, the larger is the number of cells in *P_s_* that are concentrated around *x* 1. In the limit where the DNA synthesis velocity is high, i.e. 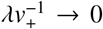, the distribution 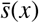 flattens and cells are equally spread along the *P _s_* phase.

We remark that while such balanced exponential growth is difficult to observe *in vivo*, where non-linear effects due to competition affecting cell proliferation, it can be observed experimentally when cells are grown in a nutrient-rich environment in the absence of contact inhibition [8, 54].

## 5. Cell-cycle progression in a dynamic environment

Having discussed the model predictions for a well-oxygenated environment, we now investigate the behaviour it exhibits under radio-biological hypoxia (RH). We start by comparing how continuous (*E*_1_) and cyclic RH (*E*_2_) affect RKO cells originally in a regime of exponential growth (i.e., the phase stationary solution computed in §4). To replicate the oxygen dynamics in the experiments from [5], we use the following functional form for the oxygen levels *c* = *c*(*t*) at time *t* > 0:

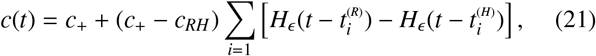

where *H*_*ϵ*_ is defined in Eq. (8c), 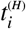 and 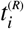 are the times at which oxygen levels decrease and increase across the threshold *c* = *c*\*, respectively. By fixing 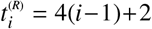 and 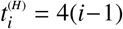, *c_RH_* ≈ 0.1% oxygen and *c*_+_ = 2% oxygen, we can reproduce the 2hr+2hr cycle corresponding to experiment *E*_2_ in Fig. 2. Fixing 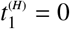 and 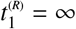, we obtain the constant RH (*E*_1_) conditions with *c* < *c*\* for *t* > 0.

### Initial conditions

Under standard culture conditions, cells *in vitro* are typically in a regime of balanced growth. We therefore initiate our simulations by assuming that cells are growing according to Eqs. (20). Recalling that we have re-scaled the model so that *N*(0) = 1, we have the following initial conditions:

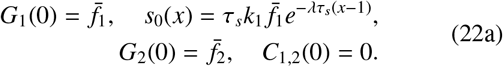

Finally, to complete Eqs. (14), we must specify the function τ_0_ (see §3.1). Since we assume DNA is synthesised at a constant rate *v*_+_ for *t* < 0, it is straightforward to show that cells with DNA content *x* at time *t* = 0 have spent a period 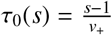 in the S phase. Using Eq. (12), we have that

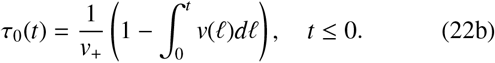

### 5.1. Numerical results

We start by considering scenario *E*_1_ where cells are exposed to constant RH for about 15 hours (see Fig. 9a and pink curve in Fig. 10). As mentioned in §2, at longer times, cell start dying and our model stops being valid, therefore we run simulations only up to this time. Fig. 10(a.1) shows that *f*_1_(*t*) rapidly increases in the first 5 hours, while it appears to settle to a value of ≈ 50% at longer times. This suggests that cells tend to accumulate in the G1 phase. By contrast, both *f_s_*(*t*) and *f*_2_(*t*) decrease. While *f_s_*(*t*) decreases monotonically over time, the decrease in *f*_2_(*t*) is delayed by a couple of hours, during which time its value remains approximately constant. Focussing now on the evolution of the model variables (see Fig. 10(a.2)), we see that the number of cells in the *C*_1_ compartment increases monotonically, but the rate of increase tends to slow after about 10 hours. By contrast, *G*_1_(*t*) slightly increases in the first few hours (≈ 4 hours) while it decreases rapidly at later times. Similarly, the *G*_2_ compartment starts to empty only after a couple of hours from the beginning of the simulation. As the velocity *v* decreases (see Fig.10(b)), the flux of cells out of the S phase (see Eqs. (17)) also decreases, contributing to the reduction in *G*_2_(*t*). Even though τ_*S*_ quickly increases above the threshold *T* ≈ 17 (see Fig.10(d)), cells do not accumulate in *C*_2_. They instead remain trapped in the S phase due to the reduction in the rate of DNA synthesis. As expected, given the trend in *G*_2_(*t*), the overall proliferation rate decreases monotonically (see Fig.10(a)), driving the gradual flattening of the population growth curve *N*(*t*) (see Fig.10(c)).

**Figure 9:**
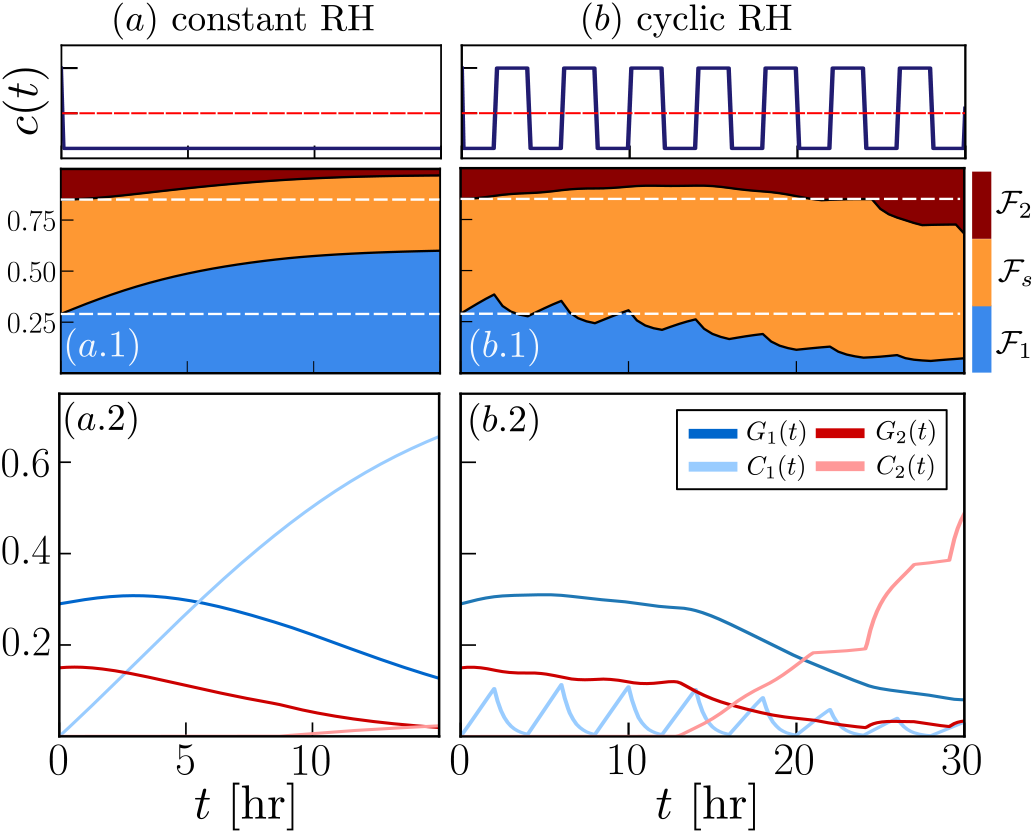
Numerical simulations of cell-cycle evolution as predicted by Eqs. (17) and (21)–(22) in constant RH (a) and cyclic RH (b). The top row illustrates the evolution of the oxygen levels *c* in the two simulations. The cellcycle dynamics are shown in the middle row panels where we plot the evolution of cell fractions over time in the form of a cumulative diagram. The white lines indicate the composition at time *t* = 0, i.e., the phase stationary solution 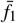, 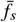 and 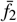. Panels (a.2) and (b.2) presents the predicted evolution of cell numbers in the different model compartments: *G*_1_(*t*), *C*_1_(*t*), *G*_2_(*t*) and *C*_2_(*t*). The parameters *k*_1_, *v*_±_ and *k*_2_ are as in Table 1 while *K*_2_ = 0 and the remaining parameter values are the estimated mean parameter values for model 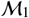 reported in Table C.9.

**Figure 10:**
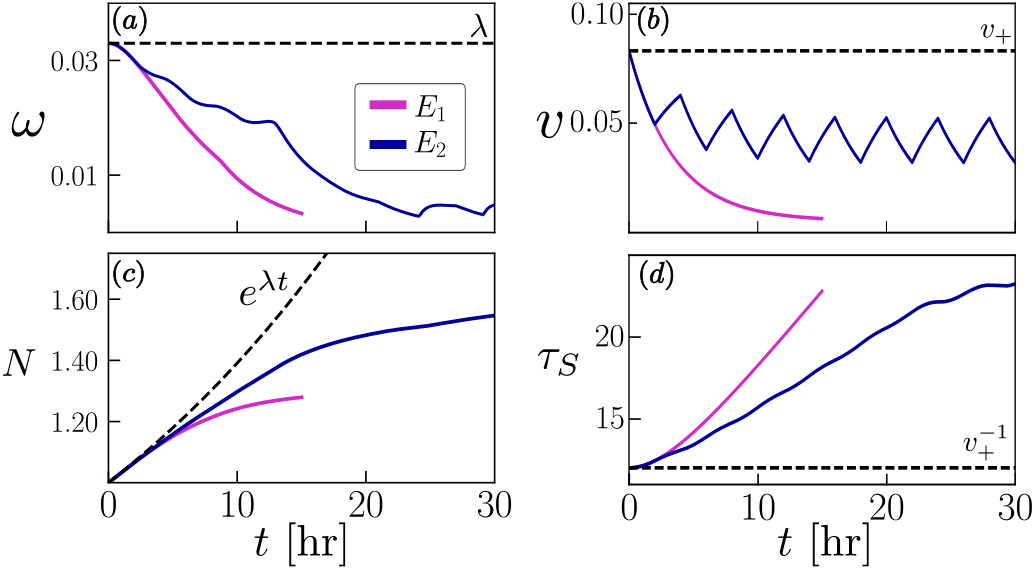
We compare the predicted population dynamics for the two simulations in Fig. 9: (a) population proliferation rate *ω* as defined by Eq. (4), (b) the total cell number *N*, (c) the DNA synthesis velocity *v* and (d) the duration of the S phase τ_*S*_. The parameters are chosen as in Fig. 9.

In the case of cyclic RH (*E*_2_), the dynamics are quite different. As shown in Fig. 9(b), while initially *f_s_*(*t*) increases and *f*_2_(*t*) decreases, the opposite occurs at later times (*t* ≈ 25 hr) when *f*_2_(*t*) increases while *f*_1_(*t*) decreases. Overall, the fraction *f*_1_(*t*) decreases, albeit non-monotonically, so that, at the end of the simulation, its value is almost negligible (i.e., 0 < *f*_1_ ≪ 1). Looking at the evolution of the number of cells in each compartment illustrated in Fig. 9(b.2), we see that cells transiently accumulate in the *C*_1_ compartment during exposure to RH (light blue curve) and resume cycling during re-oxygenation. The evolution of *G*_2_ and *G*_1_ is qualitatively similar; both remain approximately constant up to *t* ≈ 15 hr, after which time they start to decrease. At the same time, the *C*_2_ checkpoint is activated (Fig. 10(d) shows that τ_*S*_ (*t*) ≈ *T* = 17.03 hr so that *P* ≈ *p*/2 for *t* ≈ 15 hr). Since the *C*_2_ checkpoint prevents cells from replicating, its activation results in a rapid decrease in the population proliferation rate *ω* (see the blue curve in Fig. 10(a)-(b)). Under cyclic conditions, the rate of DNA synthesis falls below *v*_+_ but remains well above its minimum value *v*_−_ ≈ 5 × 10^−3^. Despite the marked fluctuations in the velocity *v*, τ_*S*_ increases almost steadily (albeit at a lower rate than for *E*_1_) until it plateaus at a maximum value of ≈ 22 hours.

As shown in Fig. 11, constant and cyclic RH also affect the evolution of the distribution *s*(*x*, *t*). Under constant RH (*E*_1_, see pink curves), cells tend to accumulate near *x* ≈ 1. Due to the low rate of DNA synthesis, the profile appears approximately stationary. In particular, comparison of the pink curves in panels (b) and (c) suggests that the discontinuity in the profile has not moved significantly. While f or *E* _1_, *s*(*x*, *t*) i s characterised by a single slow-moving front, for *E*_2_ (see blue curves) the front is followed by a series of asymmetric spikes which propagates along the *x*-axis with a faster velocity *v* (see Fig. 9(e)). Each spike corresponds to cells in the *C*_1_ compartment quickly re-entering the cell-cycle at the S phase after re-oxygenation; these cells remain highly synchronised as they proceed through the S phase. Since there is no re-oxygenation in *E*_1_, spikes are not observed. Focusing on the blue curve in Fig. 11(c), and moving from left (*x* = 1) to right (*x* = 2), the peak value decreases (as the spikes become wider). However, at later times (see Fig. 11(e)), the left-most spikes have lower peaks due to the depletion of cells in the *P*_1_ population (see Fig. 9).

**Figure 11:**
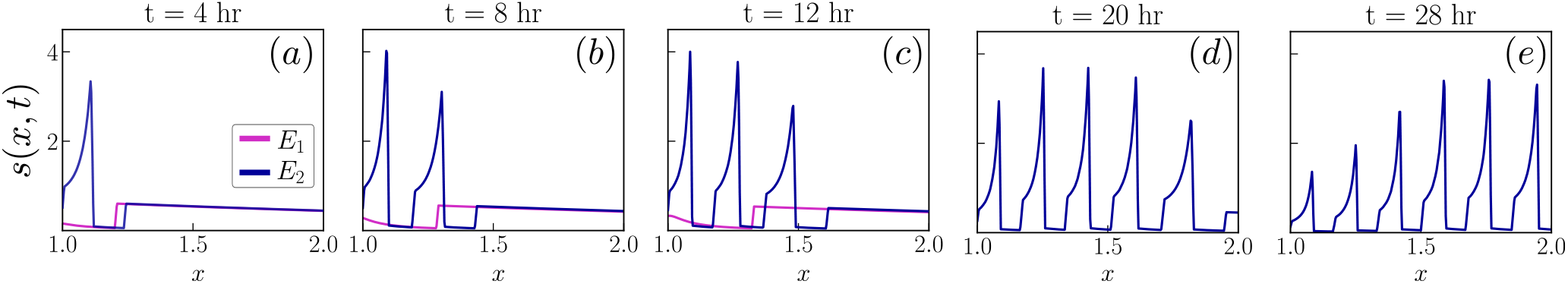
Evolution of the DNA distribution *s*(*x*, *t*) computed by numerically solving Eq. (9) for the two cases in Fig. 9: acute RH (pink curve-*E*_1_) and cyclic RH (blue curve-*E*_2_). The parameters are chosen as in Fig. 9.

Overall, our results suggest that, while both constant and cyclic RH lead to inhibition of proliferation, the mechanisms driving arrest are distinct. In the first case, cells arrest in the *C*_1_ compartment and DNA synthesis is almost completely inhibited. On the other hand, under cyclic hypoxia (*E*_2_), DNA synthesis proceeds, albeit at a lower rate. This leads to an increase in τ_*S*_ and cell accumulation in the S phase. Despite being able to complete the S phase, cells later arrest in the *C*_2_ checkpoint. This, however, is evident only at long times (≈ 24 hr), when we see a large accumulation of diploid cells. Our findings are in line with the experimental data in Fig. 2 (which are taken from [5]), indicating that our model can capture the experimentally observed cell-cycle dynamics.

Next, we use our model to investigate modes of cyclic RH not yet tested in the laboratory. For example, in Fig. 12, we fix the length of the RH phase to 2 hours and compare the growth curves for different periods of reoxygenation. We find that when the reoxygenation periods are significantly longer than the time cells spend in RH, cells have time to recover and continue proliferating, albeit at a lower rate. As shown in Fig. 13, even when cyclic RH does not result in inhibition of proliferation (i.e., cycles 2+6 and 2+8 in Fig. 12), it can still affect the cell-cycle distribution when compared to the predictions for the phase stationary solution (represented by the dashed line). For the example in Fig 13(a), cyclic RH eventually leads to *f_s_*(*t*) being above its initial value 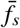. This is relevant when thinking about cell-cycle specific t reatment. S ince cyclic R H changes the distribution of cells along the cycle, it can impair or favour treatment efficacy. For example, we know that cells in different phases of the cell-cycle have different responses to radiotherapy [48]. In this case, cells in the G2/M phase have been shown to be the most sensitive to radiotherapy [48]. Referring to Fig. 13(a), we note that persistent exposure to cyclic hypoxia biases the cell-cycle distribution to the S phase. As such, it could decrease the overall sensitivity of cells to RT, even during reoxygenation, when oxygen levels do not directly increase cell radio-resistance. When considering longer re-oxygenation periods, such as in Fig. 13(b), we observe larger fluctuations in the cell fractions. Towards the end of the re-oxygenation phase, *f*_2_ is just above the value 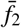. This suggests that applying treatment at this time could improve treatment efficacy. However, if treatment is not timed accurately and it is applied when *f*_2_ is at its minimum or when cells are in RH, then cyclic hypoxia could instead favour radio-resistance. This highlights the possible use of mathematical models to predict cell-cycle dynamics and how this can affect treatment outcomes when testing protocols *in vitro* accounting also for other mechanisms (such as oxygen) that can affect treatment outcomes. In order to achieve this, robust calibration of the model to experimental data is needed.

**Figure 12:**
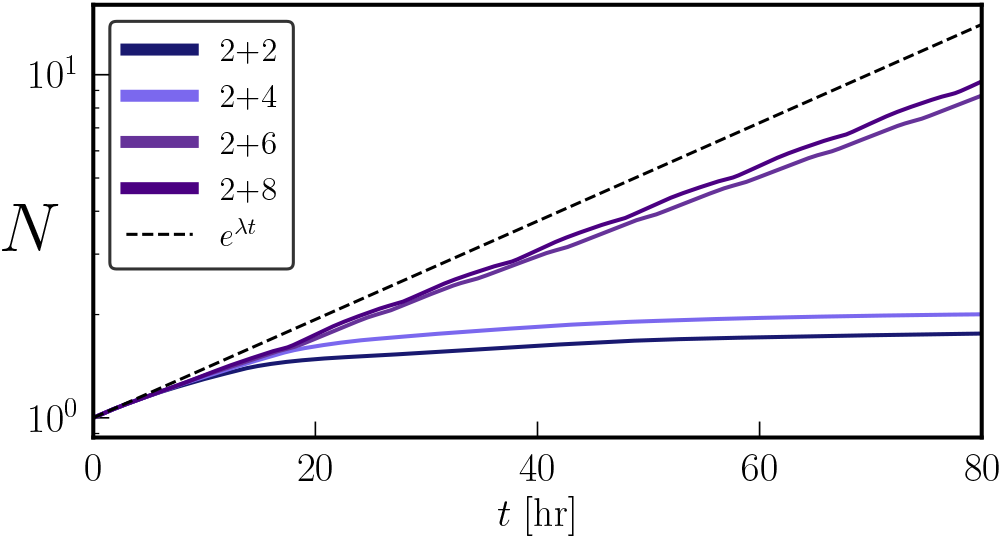
Evolution of the total cell number *N*(*t*) for different oxygen cycles as predicted by Eqs. (17). While the time spent in RH is constant for all experiments (i.e., 2 hr), we change the length of the reoxygenation phase (from 2 hr as in Fig. 9(b) to 8 hr). We also include, as a control case, the PSS (black dotted line). Parameters are chosen as in Fig. 9.

**Figure 13:**
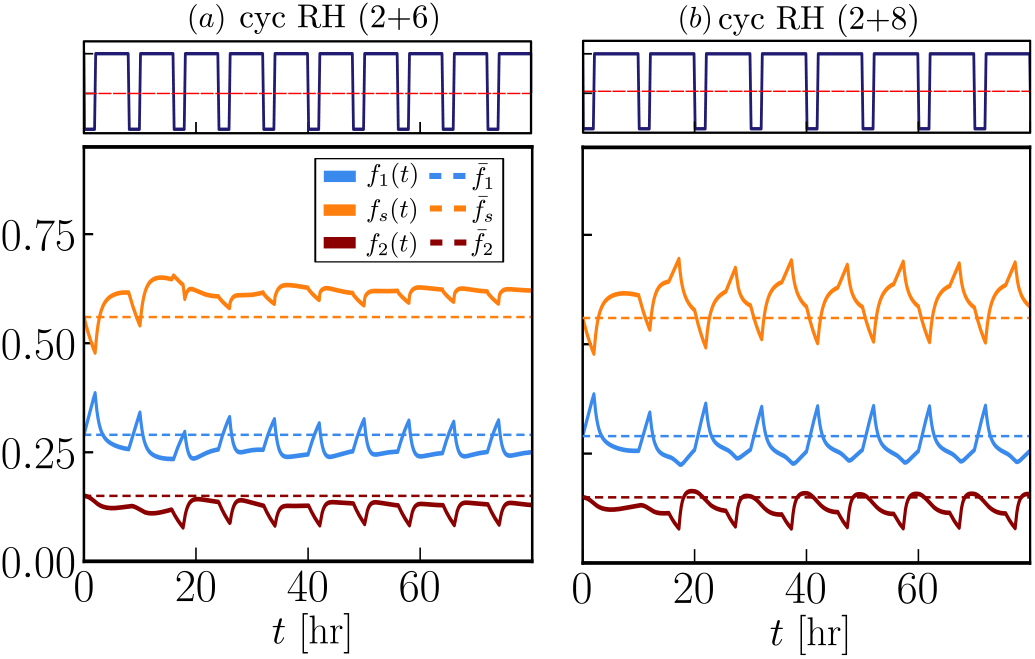
Evolution of the phase fraction *f_m_* (see solid lines) with *m* ∈ {1, *s*, 2} as predicted by the model for two different cyclic protocols in Fig. 12: (a) 2 hr (< 0.2%) + 6 hr (2%) or (b) 2 hr (< 0.2%) + 8 hr (2%). We compare the dynamics in cyclic RH with the PSS 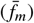 indicated by the dotted line. The parameters are chosen as in Fig. 9.

### 5.2. A class of models: comparison of different modelling assumptions

The model defined by Eqs. (17) is complex in its response to variable oxygen levels. Based on experimental evidence from [5, 27, 49], we argued in §2 that three distinct mechanisms play a key role in determining cell-cycle evolution in cycling RH. While we have shown that the model agrees qualitatively with the experimental data, it is unclear whether all three mechanisms are necessary to recapitulate the data. Further, in §5, we fixed *K* _2_ = 0 based on experimental observations; now we want to test whether this assumption is justified, based on the data available. To answer these questions, we construct a class of models, 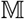 obtained by systematically reducing the complexity of the full model. A list of the models considered is presented in Table 3. While all models reduce to Eqs. (19) under normoxia, they differ under RH. The alternative models are derived from the full model, named 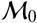, by setting 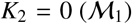 and fixing either 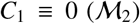 or 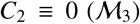. Here models 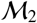 and 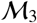 test whether the checkpoint compartments are needed to describe the data. The last two rows of Table 3 list the unknown model parameters associated with each model; while all models share the parameters ***θ***, they vary in the number of parameters ***l***^(*k*)^ associated with RH. In particular, the number of unknown parameters associated with model 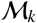 decreases as *k* increases (i.e., as the model complexity reduces). As we will detail in §6, we calibrate our models to the data in Fig. 2, and, in Fig. 14 and Fig. 15, we compare the resulting fits.

**Figure 14:**
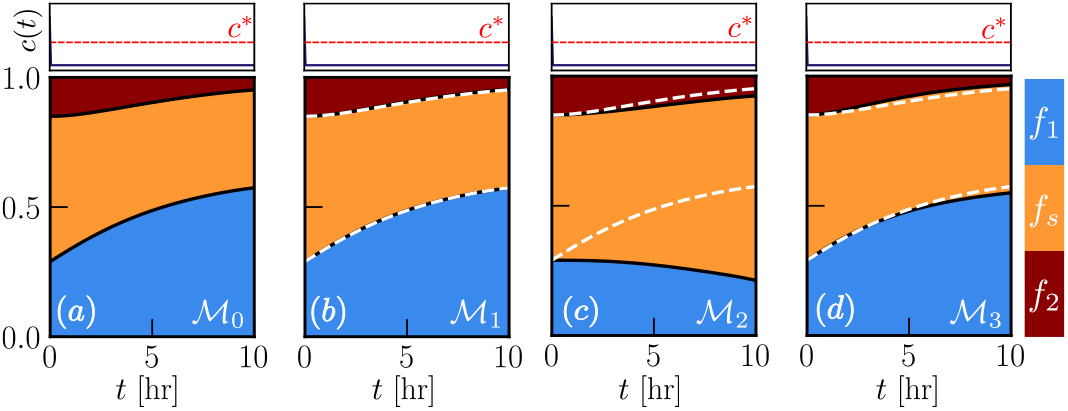
Comparison of the cell-cycle dynamics predicted by solving numerically the models listed in Table 3 for constant RH (scenario *E*_1_). We here use the best fit obtained for each model. The white dotted lines in the panels indicate the evolution predicted by model 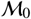 (see first panel) here used as a reference. The parameters *k*_1_, *v*_±_ and *k*_2_ are as in Table 1 while the oxygen-dependent parameters ***l*** are taken to be the estimated mean values reported in Table C.9.

**Figure 15:**
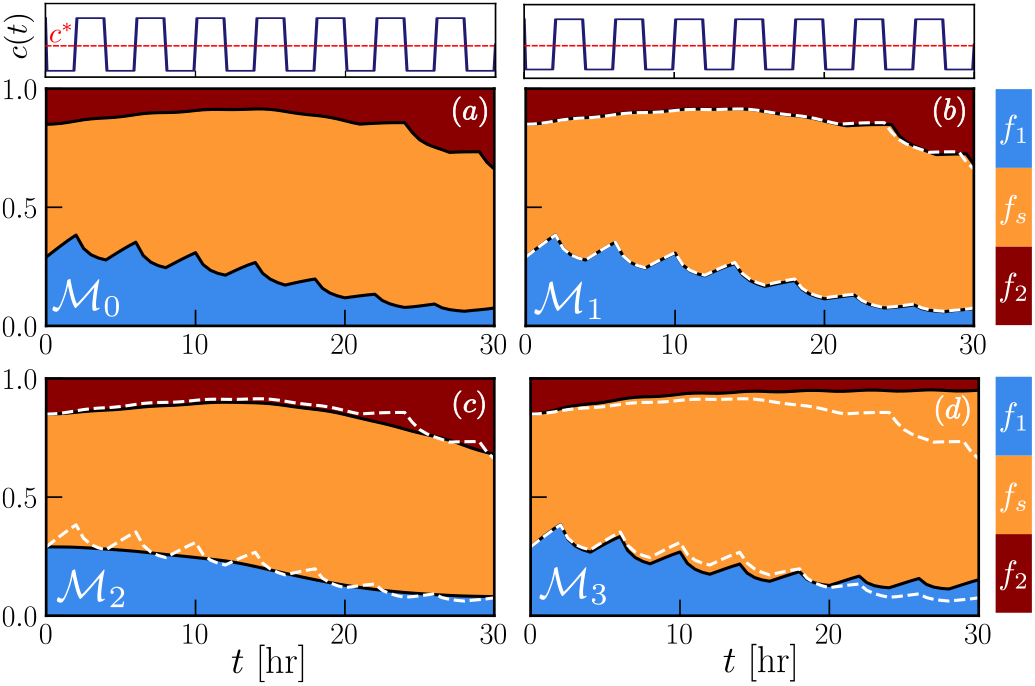
Comparison of the cell-cycle dynamics predicted by solving numerically the models listed in Table 3 for cyclic RH (scenario *E*_2_). We here use the characteristic fits obtained for each model (see Table in Appendix C). The white dotted lines in the panels indicate the evolution predicted by model 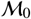 (see first panel) here used as a reference. Parameters are chosen as in Fig. 14.

**Table 3:**
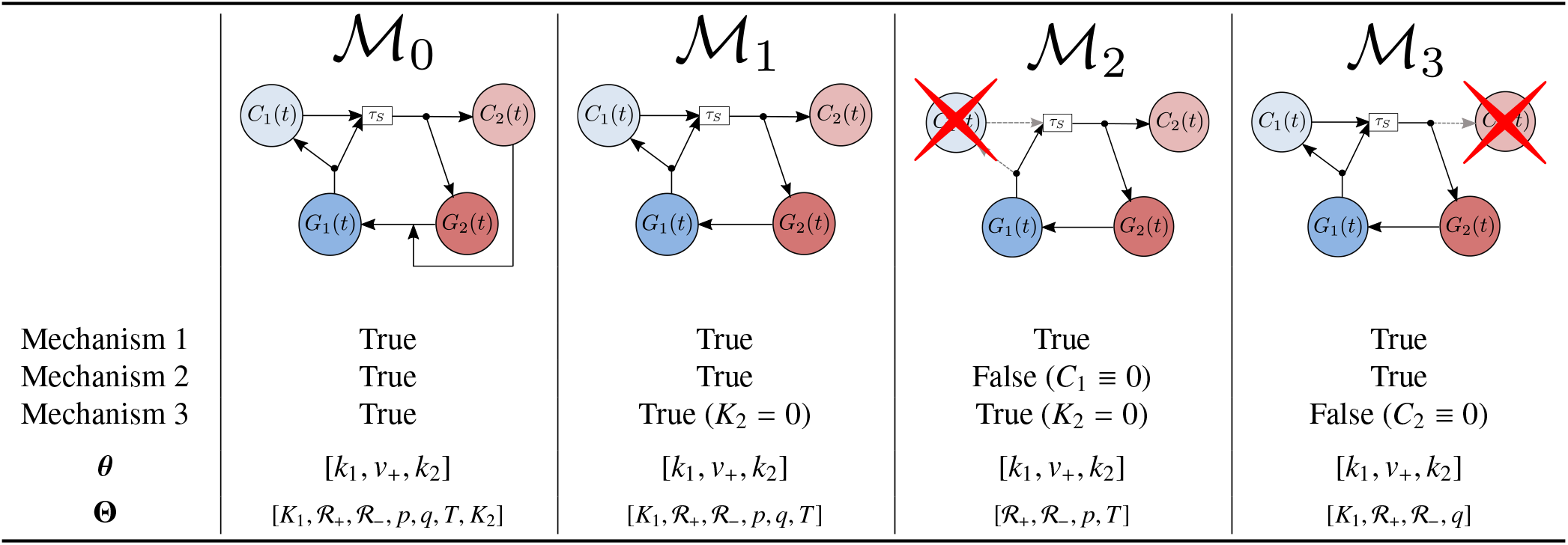
Schematics showing the biological mechanisms (see §3 for the explanation of the mechanisms (1)-(3)) included in each model 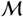 belonging to the class of models 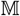. We also indicate the list of unknown parameter sets ***θ*** and **Θ** associated to each model 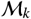 where the first set determines the phase stationary solution, while the second set plays a role in the oxygen dependent mechanisms (Mechanism 1)-(Mechanism 3). The full model, 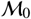, has the largest number of parameters (i.e. higher complexity) but it accounts for all the mechanisms we expect to play a role in cell-cycle dynamics under cyclic hypoxia.

| | $\mathcal{M}_0$ | $\mathcal{M}_1$ | $\mathcal{M}_2$ | $\mathcal{M}_3$ |
| --- | --- | --- | --- | --- |
| Mechanism 1 | True | True | True | True |
| Mechanism 2 | True | True | False ($C_1 \equiv 0$) | True |
| Mechanism 3 | True | True ($K_2 = 0$) | True ($K_2 = 0$) | False ($C_2 \equiv 0$) |
| $\theta$ | $[k_1, v_+, k_2]$ | $[k_1, v_+, k_2]$ | $[k_1, v_+, k_2]$ | $[k_1, v_+, k_2]$ |
| $\Theta$ | $[K_1, \mathcal{R}_+, \mathcal{R}_-, p, q, T, K_2]$ | $[K_1, \mathcal{R}_+, \mathcal{R}_-, p, q, T]$ | $[\mathcal{R}_+, \mathcal{R}_-, p, T]$ | $[K_1, \mathcal{R}_+, \mathcal{R}_-, q]$ |

In Fig. 14, we compare predictions when cells are exposed to constant RH (*E*_1_). In this case all models, except 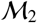, predict that *f*_1_ increases in line with the experimental observations from [5]. In contrast, model 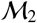 (Fig. 14(c)) is incompatible with the experimental data, predicting a monotonic decrease in *f*_1_(*t*) and cell accumulation in the S phase (observe the large increase in *f_s_*(*t*) over time). This suggests that in constant RH, the *C*_1_ checkpoint plays a key role while the *C*_2_ checkpoint can be neglected given that model 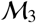 qualitatively reflects the same trends observed experimentally. However, neglecting the *C*_1_ checkpoint would not significantly impact predictions under cyclic RH. Indeed, looking at Fig. 15(c), we note that model 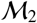 is in good agreement with model 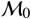 and 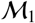 (indicated by the white dotted line). While 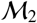 does not capture the rapid fluctuations in *f*_1_(*t*) predicted by 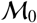, the overall trend is the same, with *f*_1_(*t*) decreasing after each cycle. Focusing on Fig. 15(d), we see good agreement between 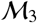 and 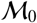 only for the first 20 hours. At later times, 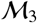 predicts values for *f*_2_ significantly lower than those predicted by 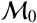 and observed experimentally (see Fig. 2). This suggests that the *C*_2_ checkpoint is needed for the model to capture the delayed accumulation of cells in the G2/M phase observed experimentally.

The results from Figs. 14 and 15 highlight the inability of models 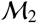 and 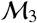 to recapitulate the experimental data. Further, the predictions from 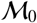 and 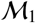 are indistinguishable, which suggests that the value of the parameter *K*_2_ does not significantly influence the dynamics. Discriminating between models 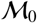 and 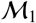 is, therefore, not straightforward and requires consideration of other metrics, in addition to how well the model fits the experimental data. These questions will be addressed in the next section where we implement *Bayesian model* selection.

## 6. Parameter fitting and model selection

So far we have presented model predictions for point estimates of the model parameter values. In this section, we explain how such estimates were obtained and investigate how the results in §5 change when we account for uncertainty in the estimates of “oxygen-dependent” parameters. We start by using the data from the balanced exponential growth experiments (*E*_0_) to determine the “oxygen-independent” parameters ***θ*** using the results from §4. As mentioned in §5.2, all models 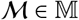 reduce to the same set of equations (Eqs. (19)) in normoxia. As such, they share the same value ***θ***. We then focus on estimating the remaining parameters **Θ**, which determine the system response to dynamic oxygen conditions. Here we will compare different modelling assumptions by applying model selection methods. We start by fitting each model 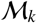 in Table 3 to experimental data from both *E*_1_ and *E*_2_ simultaneously, using Bayesian inference and Monte Carlo methods to estimate the posterior distribution, π_*ps*_, for the parameters **Θ** (more details follow). We then select the “best” candidate model by using the deviance information criterion (DIC) as an estimate of model performance and briefly discuss parameter identifiability based on posterior profiles.

### 6.1. Estimation of oxygen independent parameters

We recall from §4, that the regime of unperturbed exponential growth is characterised by the value of the constants 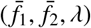. As explained further in Appendix A, these constants uniquely define the values of the parameters (*k*_1_, *k*_2_, *v*_+_) given by Eqs. (A.7). In practice, we can estimate the stationary values 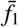 and 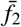 using flow cytometry data. However, additional data are needed to determine the proliferation rate λ. This parameter can be related to the population doubling time, *T_doub_* [8], which is known for most cell lines when cultured in standard media and in the absence of competition (i.e., low confluence). Indeed, it is straightforward to show that *T_doub_* = λ^−1^ ln 2.

Given that, prior to any of the experiments in [5], the cells were cultured at 21% *O*_2_, with re-plating in order to minimise any effects due to contact inhibition, we assume that initially the cells are undergoing exponential growth. We can, therefore, use the cell fractions data reported at time *t* = 0 in Fig. 2 to estimate the values of 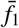 and 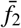. For simplicity, we suppose that median values provide a good approximation to the ‘true’ cell fractions so that 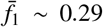 and 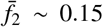. In this way, we can obtain point estimates for ***θ*** which facilitates the identification of the remaining model parameters **Θ**. In previous experiments, the doubling time of RKO cells has been estimated to be about 21 hours [65]. The corresponding parameter estimates (obtained using Eq. (A.7)) are listed in Table 4 together with estimates of the duration of each cell cycle phase (given by the inverse of the rates of *k*_1_, *k*_2_ and *v*_+_ [8]). We note that the RKO cell line is characterised by a particularly long S phase. By contrast, the durations of the G1 and G2/M phases are almost half the duration of the S phase, with G2/M being the shortest.

**Table 4:** Estimates of the cell-cycle parameters for the RKO cell line obtained using the phase stationary solution (PSS) and the experimental data from [5].

|  | transition rates<br>[1/hr] | average time in the compartment [hr] | cell fractions<br>(from [5]) |
| --- | --- | --- | --- |
| G1 | $k_1 = 0.195$ | $\tau_1 \approx k_1^{-1} \approx 5.14$ | $\bar{f}_1 = 0.29$ |
| S | $v_+ = 0.083$ | $\tau_S = v_+^{-1} \approx 12$ | $\bar{f}_s = 0.56$ |
| G2/M | $k_2 = 0.22$ | $\tau_2 \approx k_2^{-1} \approx 4.54$ | $\bar{f}_2 = 0.15$ |

### 6.2. Calibration of the candidate models to time-series flowcytometry data

The second step concerns estimation of the parameters **Θ** which are associated with the oxygen-dependent mechanisms. Here we use Bayesian inference techniques [41], which allow us to account for measurement errors and to assess uncertainty in the parameter estimates.

Given the small amount of data available, we calibrate the model by pooling the data from all of the experiments in constant RH (E1) and cyclic RH (E2). We therefore postpone model validation until more data will be available. In Fig. 2, for each time point, we reported the mean and standard deviation for multiple repeats of the experiment; however, for the estimation of **Θ** we consider individual experimental measurements, instead of summary statistics. The complete data set can be found in Appendix B. We denote by ***F***^(*i, j*)^ the i-th measurement performed at time *t*^(*i, j*)^ during experiment *E*_*j*_ with *j* ∈ {1, 2}. Given that *F* _1_, *F* _*s*_ and *F*_2_ are not independent (recall *F* _s_ = 1 − *F* _1_ − *F* _2_), we define the values of **F** =[*F* _1_, *F* _2_] ∈ ℝ^2^. we collect the data in the set 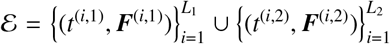, where *L*_1_ = 4 and *L*_2_ = 14 are the number of measurements in sets *E*_1_ and *E*_2_, respectively.

### The error model

Details of the error between the experimental measurements and the predictions of the mathematical model 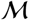, with model parameters **Θ**, are captured in the *likelihood* function 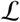. We assume the measurement errors are additive and independent of the oxygen protocol *c* = *c*(*t*) used in the *in vitro* experiments. We further suppose that the difference between each measurement and model prediction 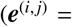 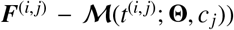 to be independent and normally distributed, with covariance matrix **Σ**, i.e., 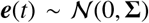; therefore 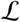 has the form:

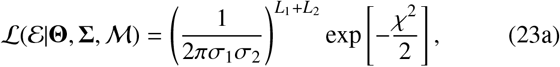

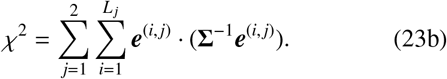

Here the covariance matrix **Σ** is considered to be constant in time and diagonal (i.e., 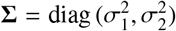), as we assume independence between errors in the fractions *F*_1_ and *F*_2_. Instead of specifying the values of σ_1_ and σ_2_, we treat them as unknown parameters that are learnt from the data. Since the cell fractions are normalised, we assume that σ_1_ and σ_2_ are uniformly distributed in [0, 1] 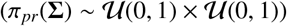.

### Calibration and model selection

In brief, given a mathematical model 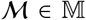, the main goal of Bayesian inference is to sample, via Monte Carlo methods, from the posterior probability distribution for the model (**Θ**) and error parameters (**Σ**) conditioned to the data 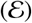 (i.e., 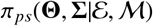). Using Bayes’ Theorem, the posterior distribution can be expressed in terms of the *likelihood* function 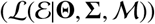 and the prior distributions 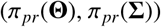:

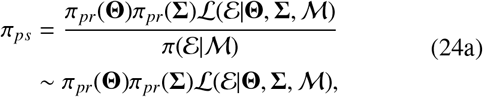

where 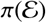 is a normalising factor, π_*pr*_ is the prior knowledge on parameter values, and 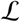 is as defined in (23b).

In the absence of prior information about the values of model parameters **Θ**, we assume that each component of the vector **Θ** (i.e., **Θ**_*i*_ for *i* = 1, … , κ), is uniformly distributed on the intervals 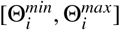 where the extremes of the intervals are taken from Table 2; hence π_*pr*_(**Θ**) is given by:

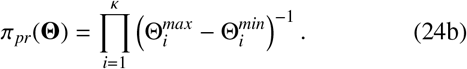

Due to the dimensions of our parameter space, it is not possible to compute Eqs. (24a) analytically. Instead we rely on Markov Chain Monte Carlo (MCMC) methods to estimate π_*ps*_ using the python package PINTS (Probabilistic Inference on Noisy Time-Series) for Bayesian inference [19]. More details on the numerical technique are included in Appendix C.

For model selection among the class of models 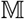, we compute the deviance information criterion (DIC) [28, 41]. The DIC represents a trade-off between model complexity (as measured by the over-fitting bias *k_DIC_*) and model accuracy (as measured by the likelihood). Given a model 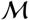, the DIC is defined as follows [41]:

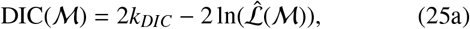

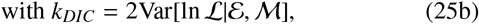

where 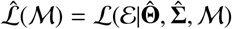 is the likelihood value estimated at the expected value of the parameters, (i.e., 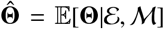 and 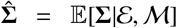) and Var is the variance of the log-likelihood, ln 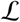, here approximated via sampling from the estimated posterior. We note that the DIC penalises models with large values of *k_DIC_*, to account for the fact that complex models are more likely to fit data well. At the same time, more complex models tend to have more parameters which can lead to higher posterior uncertainty if the model is too complex for the data (i.e., it is over-fitted) [41]. This results in a larger variability in the expected log-likelihood (i.e., larger *k_DIC_*). In other words, the term *k_DIC_* in Eq. (25a) corrects for over-fitting. When comparing models, we are interested in the relative value of the DIC, and favour the model with the smaller DIC.

For a model 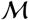, predictive posterior estimates are obtained by sampling 800 parameter sets, (**Θ**^(*i*)^, **Σ**^(*i*)^), from the estimated posterior π_*ps*_. For each set, we run the model 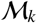 forwards to generate 800 predictive curves for each phase fraction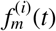 with *m* ∈ {1, 2} and *i* = 1, … , 800. This gives posterior distributions for the *“true”* cell fraction. To obtain posterior distributions for the *measured* cell fraction 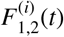, we add to the simulated cell fractions 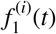 and 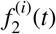 the corresponding measurement errors 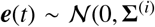). We then estimate 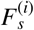 as 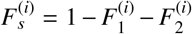 and 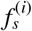 as 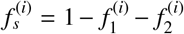. At each time point, we compute the mean of the 800 predictive curves (for either the “true” or measured fractions) and the corresponding 68%- and 95%-confidence intervals. For the plots in §5 and §5.2, we used the expected values 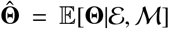 as representative of characteristic model fits. Additional results on the estimated posterior distributions can be found in Appendix C.

#### 6.2.1. Numerical results

The estimated DICs for the models 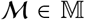 are reported in Table 5. Based on these estimates, model 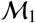 has the lowest DIC and is, therefore, the “best” model in the class 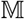. As expected from the results presented in §5.2 (see Fig. 14 and 15), 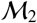 and 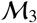 have the worst performance. In this case, the difference in the estimated DICs is rooted in the value of 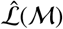, which is markedly reduced for 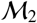 and 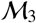, suggesting that the two models fail to fit the experimental data. We conclude that including the *C*_1_ and *C*_2_ compartments (*i.e.* cell-cycle arrest in the G1 and G2 phases) is necessary for our model to reproduce the experimental cell-cycle dynamics. However, we can not exclude the possibility that other mechanisms (i.e., modelling assumptions) not considered here, might also explain the data.

**Table 5:** Comparison based on the deviation information criterion (DIC) of the models in class 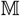 that were introduced in §5.2. The last column indicates the relative *DIC* score with respect to model 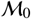, i.e., 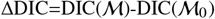.

| | $k_{DIC}$ | $\ln(\hat{\mathcal{L}}(\mathcal{M}))$ | DIC | $\Delta\text{DIC}$ |
| --- | --- | --- | --- | --- |
| $\mathcal{M}_0$ | 6.10 | 65.73 | -119.27 | 0 |
| $\mathcal{M}_1$ | 5.13 | 65.33 | -120.41 | -1.73 |
| $\mathcal{M}_2$ | 6.04 | 43.83 | -75.58 | 43.62 |
| $\mathcal{M}_3$ | 4.25 | 45.84 | -83.18 | 37.35 |

Focusing now on 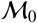, we see that its performance is similar to that of 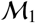, whose DIC is only slightly smaller. While the two models yield similar values of 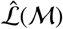, the larger estimated value of *k_DIC_* for 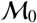 is the main source of the discrepancy. This suggests that the additional complexity due to the introduction of the parameter *K*_2_ is not balanced by a sufficient improvement in the description of the data. This interpretation is confirmed by the profile of the marginal posterior for *K*_2_ (see Fig. 16(a)). The latter is approximately uniform, suggesting that its value can not be identified, given the available data. As shown in Fig. 16(b), the posterior predictions for *f*_2_ have larger confi-dence intervals at longer times. This suggests that experiments run over longer time periods could improve the estimates of parameter *K*_2_. In the absence of such data, we view model 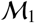 as the best candidate.

**Figure 16:**
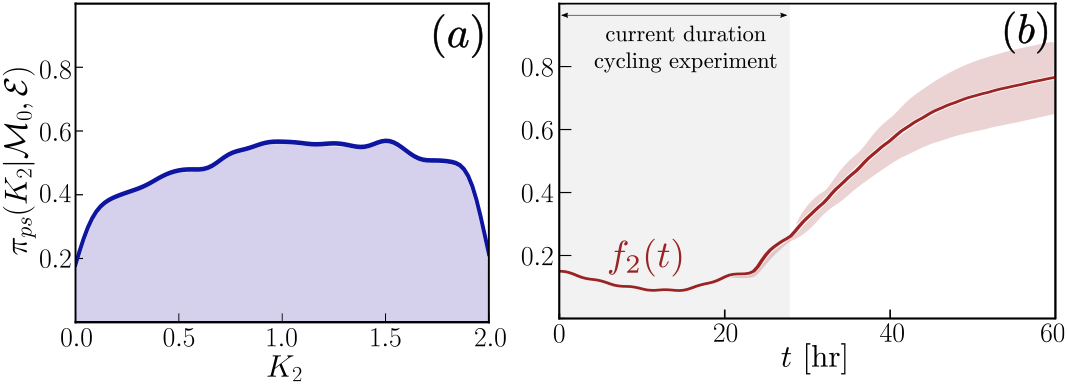
Analysis of the fitting of model 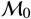: (a) marginal posterior distribu-tion for parameter *K*_2_; (b) posterior prediction of the time evolution of *f*_2_, i.e., the fraction of cells in the *P*_2_ phase. The mean (red curve) and 68%-confidence intervals (red shaded area) are computed as outlined in the section on “Calibration and model selection”.

### Candidate model M_1_ is in good agreement with the experimental data

Our analysis shows that model 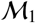 gives the best fit to the experimental data. Referring to the posterior predictions in Fig. 17, we note that 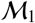 captures the experimental data from both experiments *E*_1_ and *E*_2_, with all experimental data points falling within the 95%-confidence interval of the posterior predictions for the “measured” fractions, *F*_1_, *F_s_* and *F*_2_.

**Figure 17:**
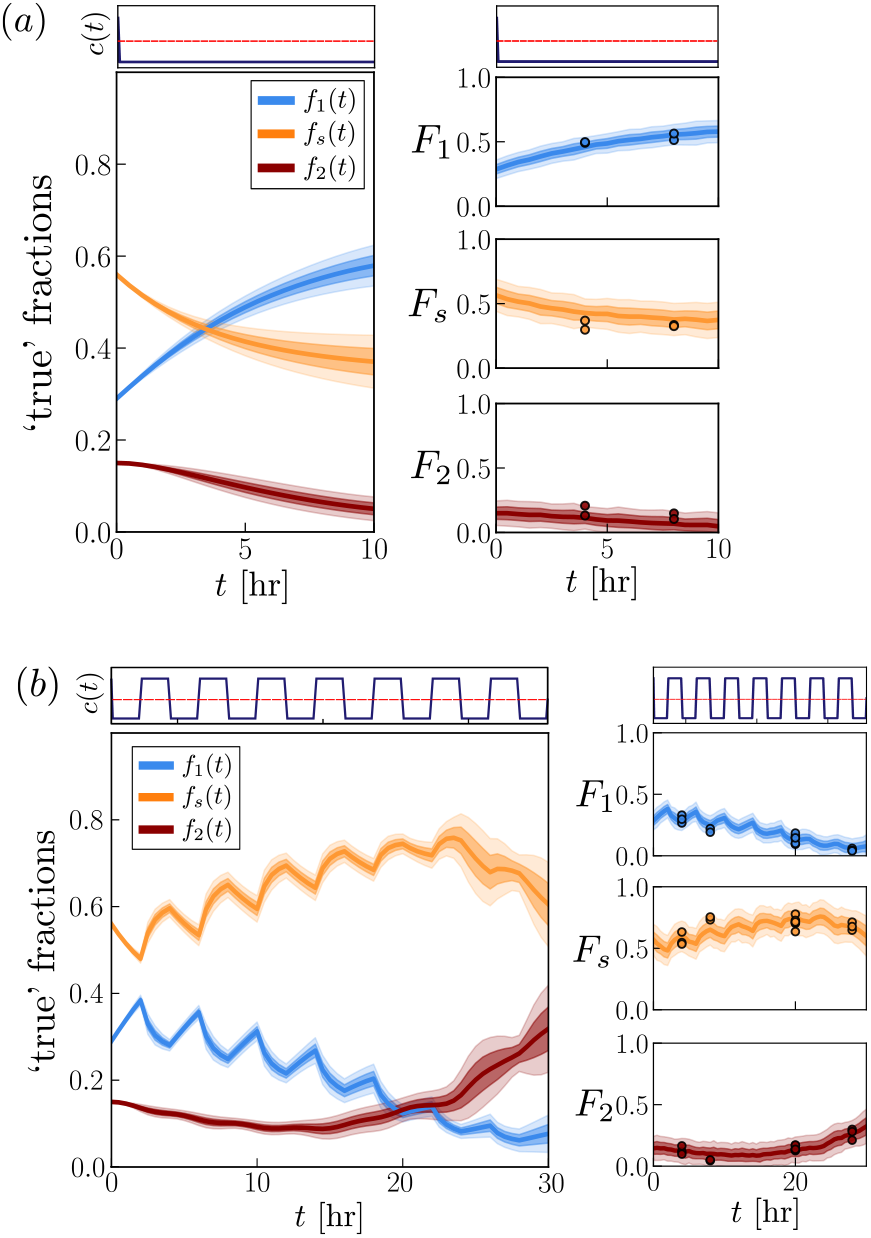
Posterior prediction distribution of the selected model 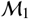 for (a) chronic RH and (b) cyclic RH. For both scenarios we plot the predicted ‘true’ cell fractions *f_m_*(*t*) and the predicted measured fractions *F_m_*(*t*) with *m* ∈ {1, *s*, 2}; the predicted evolution of *F_m_* is compared with the experimen-tal data (dots). For each model output, we plot the expected values, together with the 68%- and 95%-confidence intervals, as indicated by the dark and light shaded areas, respectively. The top panels indicate the prescribed oxygen tension *c*(*t*) associated with the two experiments.

Fig. 18 shows that 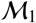 can also capture the qualitative shape of the flow cytometric distribution (*D_f_*) reported in [5], which indicates cell number over PI-fluorescence intensity (see Fig. 1). Here we estimate *D_f_* using output from the numerical solution of Eqs. (17) as follows: based on [10], we assume that a population of cells with the same DNA content *x* gives rise to a Gaussian-like flow cytometric output ( see Fig 1); consequently, we model the flow cytometric output *D_f_* as

**Figure 18:**
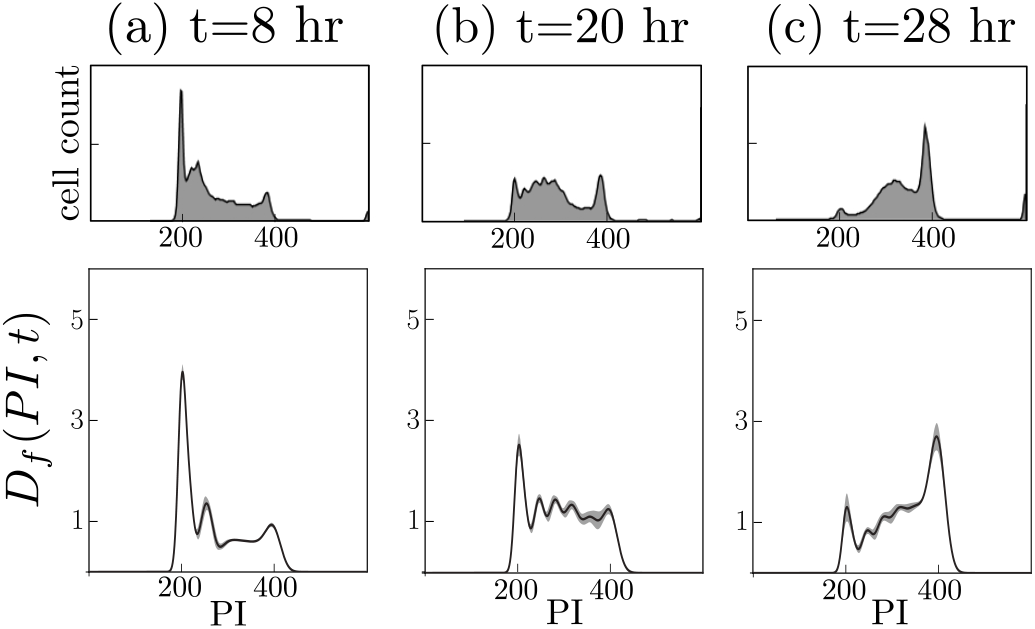
Posterior prediction for the distribution *D_f_* as defined by Eq. (26a) for cyclic RH (*E*_2_). The dark line indicates the average over 800 parameter sets sampled from the estimated posterior, while the shaded grey area indicates the 68% confidence interval. At each time point, we compare the theoretical prediction (bottom panels) with the experimentally measured distribution from Fig. 3 in Bader et al. [5] (top panels). Fluorescence readings of PI = 200 and PI = 400 correspond, respectively, to *x* ≈ 1 and *x* ≈ 2. This comparison is qualitative, as the vertical axes for the theoretical (bottom panels) and experimental (top panels) distributions are different since *D_f_* is re-scaled by the initial number of cells *N*_0_, while the experimental output is in terms of absolute cell counts.

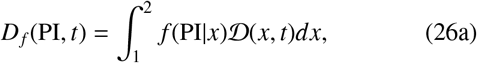

where

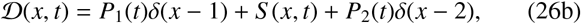

where *δ*(*x* − *y*) is the delta function, (*i.e.*, *δ* = 1 if *x* = *y* and *δ* = 0 otherwise). The term *f* (PI | *x*) in Eq. (26a) is the probability of recording a fluorescence intensity *PI* for a cell with DNA content *x* and has the form

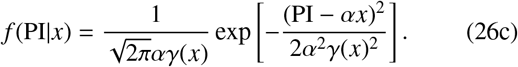

In Eq. (26c), we assume that the mean PI fluorescence is proportional to the DNA content *x*, with constant of proportionality α = 200 PI, so that *x* = 1 corresponds to a fluorescence intensity PI = 200. Following [10] again, we suppose that the variance *γ* depends linearly on *x* (*i.e.*, *γ* = (*γ*_2_ − *γ*_1_)(*x* − 1) + *γ*_1_, with *γ*_1,2_ being the variances associated with DNA contents of *x* = 1 and *x* = 2, respectively). In general, *γ*_1_ and *γ*_2_ will depend on the cell line of interest. Since here we are interested in qualitative comparisons, we set *γ*_1_ = 0.04 and *γ*_2_ = 0.08, for consistency with the estimates from [10].

While the calibration only uses information about cell fractions, we see that our theoretical estimates for *D_f_* are in good agreement with the experimental observations (see Fig. 18). However, there are some discrepancies, particularly at time *t* = 20 hr. The model predicts a higher percentage of cells in the late S phase, so that the peak corresponding to *PI* = 400 is not isolated; in contrast the experimental profile tends to flatten in the vicinity of PI ≈ 350, so that the peak at PI ≈ 400 is isolated. However, we note that the experimental observations relate to only one realisation, whereas we illustrate the expected profile over several model realisations.

### Model Identifiability

We end our discussion on model calibration by briefly considering practical identifiability of the unknown parameters. Following [23, 35], we define a parameter as practically identifiable if we can constrain its value to a reasonably small region of parameter space, *i.e.*, the posterior distribution has compact support.

Looking at the profile of the marginal posterior distributions for the different parameters (see Fig. 19), we see that the marginal posteriors have a bell-like shape, with a unique, welldefined, maximum for most parameters. However, the posterior distribution for parameter *K*_1_ tends to flatten at large values, where *K*_1_ > 1 (see Fig. 19(a)). This indicates greater uncertainty in the estimation of *K*_1_ for which we can identify only a lower, but not an upper, bound. From Eq. (8a), we note that *K*_1_ is only relevant in data from experiment *E*_2_ (i.e., when oxygen levels *c* are above the threshold *c*\*). If we consider, for example, the first time at which this happens (*i.e.*, time 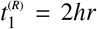), then, over the period 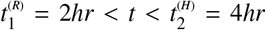, *Q*(*t*) ≈ 0 and the evolution of *C*_1_(*t*) can be determined explicitly by solving Eq. (17b) to obtain:

**Figure 19:**
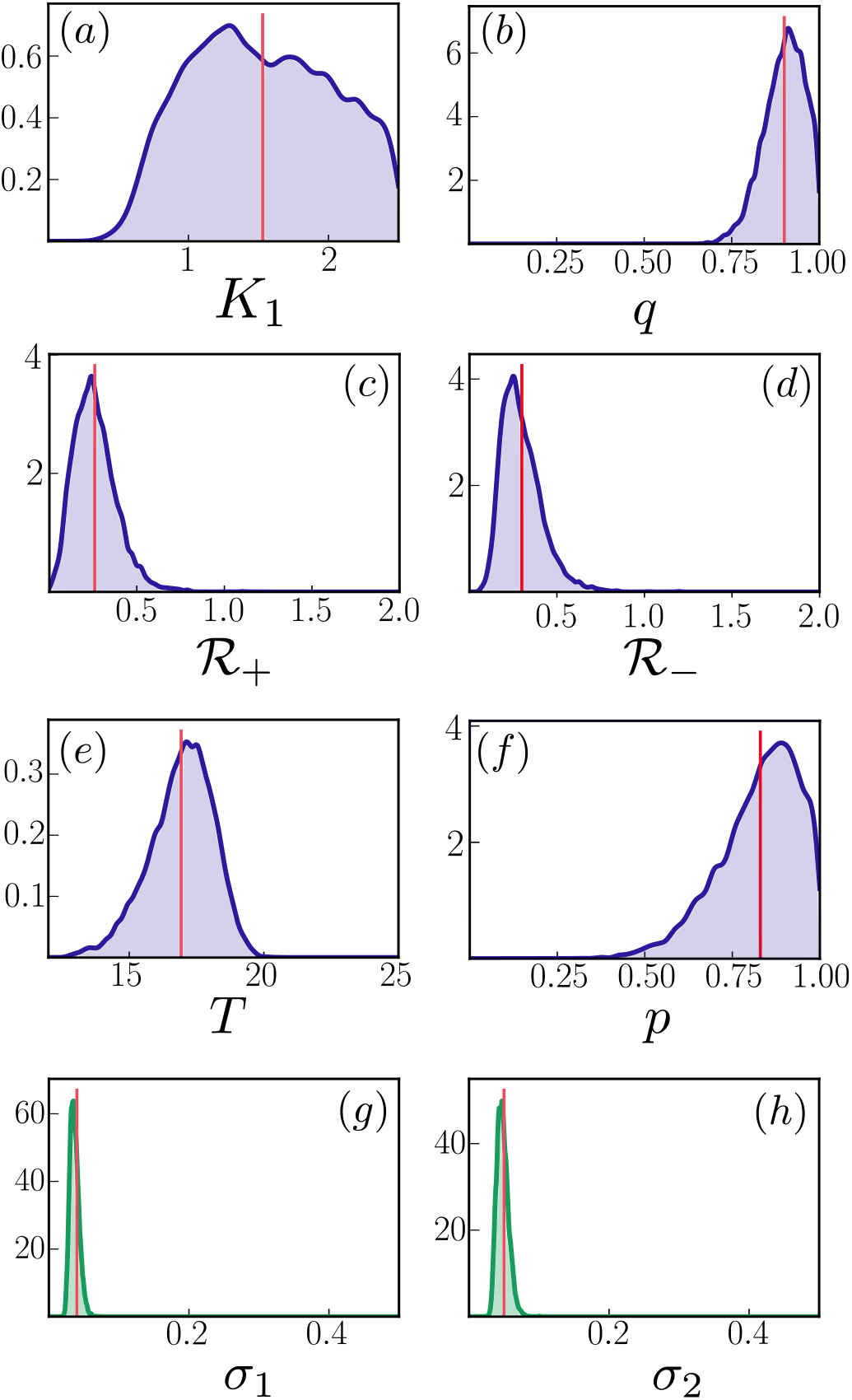
Marginal posterior distributions 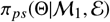 for the model parameters 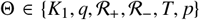 and the error parameters 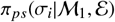 for *i* = 1, 2. The red vertical line indicates the mean value as reported in Table C.9. We see that for most parameters the posterior distribution has a compact sup-port. This is not the case for *K*_1_ (see (a)) where the large support of the posterior is a sign of practical non-identifiability.

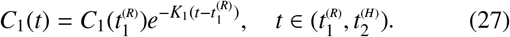

Thus, when the first measurement is taken, 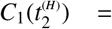 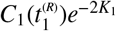. If *K*_1_ > 1, then 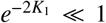 and compartment *C*_1_ rapidly empties after re-oxygenation. Therefore, unless *C*_1_ is very large, 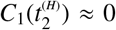, independently of the value of *K*_1_ > 1. In order to resolve this fast time-scale we would need to collect experimental data at an earlier time point, say *t*\*, for which 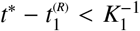; alternatively, we could choose an oxygen cycle for which a larger number of cells accumulate in the *C*_1_ compartment. This could be achieved by prolonging the period for which the cells are exposed to severe hypoxia (i.e., *c* < *c*\*).

## 7. Model predictions in the presence of uncertainty

To conclude, we use the calibrated model from §6 to make predictions on cell-cycle dynamics in different environmental conditions. We start by considering the oxygen cycles in Fig. 13 but now account for uncertainty in our parameter estimates.

For the 2hr+6hr cycle in Fig. 20(a), in the absence of uncertainty, we predicted a systematic increase in the fraction of cells in the S phase, with no activation of the *C*_2_ compartment (see Fig. 13(a)). When uncertainty is taken into account, we find large variability in model predictions at longer times (*t* > 25 hr). In particular, the 68%-confidence interval encompasses the possibility of *f_s_*(*t*) both increasing or decreasing compared to its initial value when *t* ≫ 1. Further, activation of *C*_2_ cannot be ruled out, as indicated by the value of *f*_2_(80), which ranges between 0 ⪅ *f*_2_(80) ⪅ 0.6 (see Fig. 20(a.3)). Despite the large uncertainty in the value of *f*_2_(*t*), the confidence interval on *f*_1_(*t*) remains reasonably small. We observe similar behaviour for cycles with a longer re-oxygenation phase (see 2hr+8hr cycle in Fig. 20(b)). Again, uncertainty in *f_s_* and *f*_2_ increases over time, even though it is less pronounced than in Fig. 20(a). Again, the 68%-confidence interval allows for the possibility of activation of the *C*_2_ checkpoint (see increase in *f*_2_ as *t* ≫ 1) even though this is less probable than in the scenario depicted by Fig. 20(a). For both oxygen cycles considered, the uncertainty in the cell-cycle distribution is reflected in the predictions for the number of cells *N*(*t*) (see Fig. 21). This is particularly evident in Fig. 21(a) where *N*(80), the number of cells at the final time, *t* = 80 hr, ranges between 3.5 ⪅ *N*(80) ⪅ 10.

**Figure 20:**
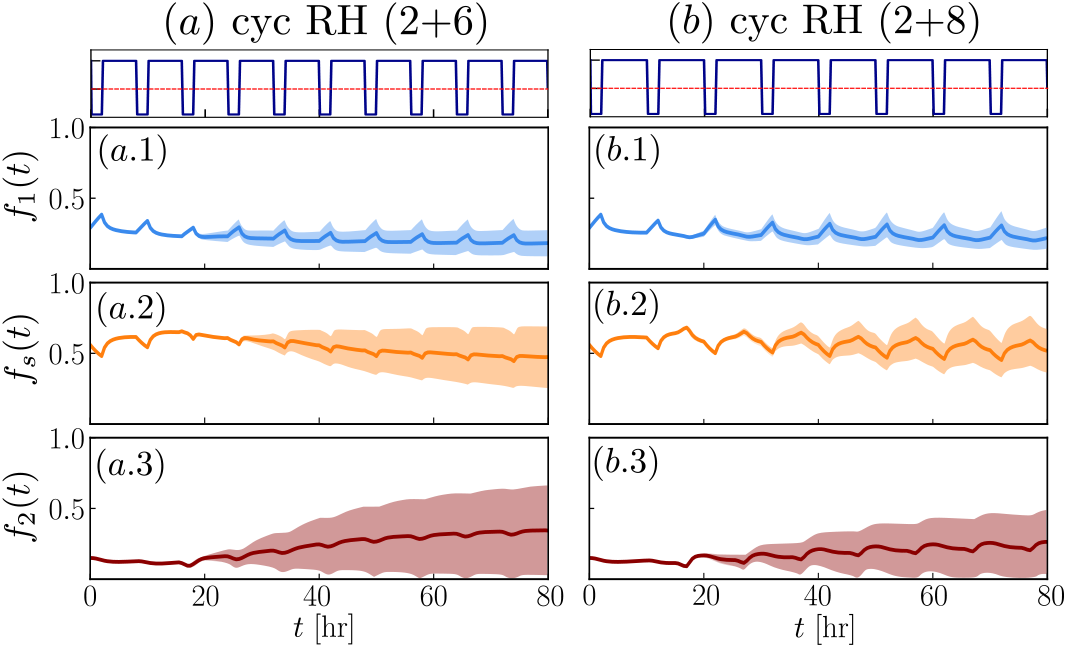
Posterior estimates for the cell fractions *f_m_* for the two cyclic protocols considered in Fig. 13: (a) cyclic RH with 2 hr in RH + 6 hr in an oxygenated environment; (b) cyclic RH with 2 hr in RH + 8 hr in an oxygenated environment. We plot mean estimates and indicate the 68%-confidence intervals by the shaded regions.

**Figure 21:**
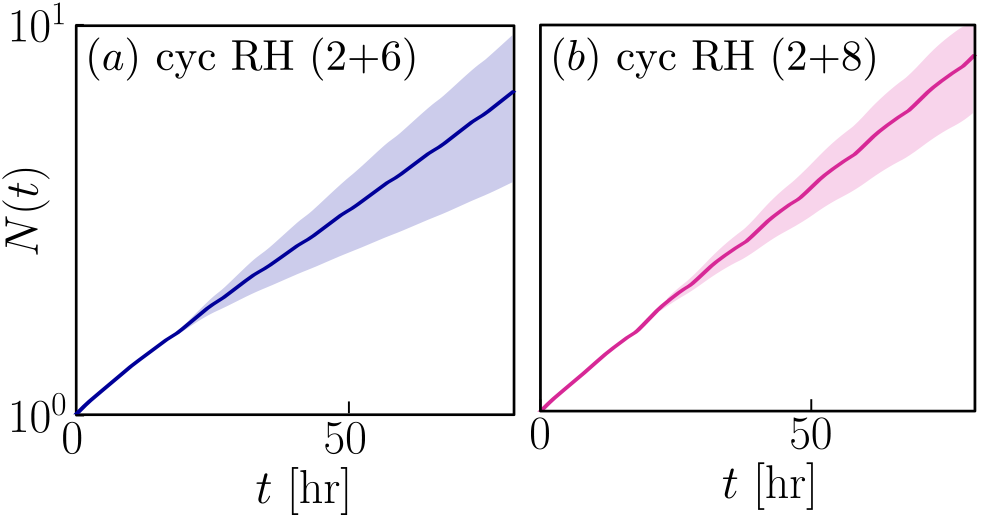
Posterior estimates for the total cell number *N*(*t*) for the two cyclic protocols in Fig. 20: (a) cyclic RH with 2 hr in RH + 6 hr in an oxygenated environment; (b) cyclic RH with 2 hr in RH + 8 hr in an oxygenated environment. We plot mean estimates and indicate the 68%-confidence intervals by the shaded regions.

Based on the results in Fig. 20, we conclude that we can use our calibrated model to predict cell-cycle dynamics on short time scales (0 < *t* < 25 hr); thereafter the increased uncertainty prevents us from making reliable predictions for the cell-cycle distribution at longer times. However, the predicted uncertainty is still informative when considering experimental design. In this case, the objective is twofold. On the one hand, we want to identify experiments that could facilitate model validation. From this point of view, we focus on times at which the uncertainty in our predictions is small (e.g., on the short time scale for experiments in Fig. 20). On the other hand, we aim also to propose experiments that could improve the accuracy of model parameter estimates. In this case, our attention focuses on scenarios where uncertainty in the model predictions is large and, therefore, new measurements can refine parameter estimates (such as for the long time dynamics in Fig. 20). From this point of view, a scenario like Fig. 20(a) is adequate since it can account for both the validation and refinement steps. Our model also suggests that, if such experiments were performed, information about the total number of cells *N*(*t*) may improve model calibration, given the large variation predicted in its value at the end of treatment.

Experimentally, it might be difficult to study scenarios for which oxygen levels change on timescales faster than two hours due to the time required for oxygen levels to equilibrate *in vitro* [50]. This, however, is not a limitation of our model. Indeed, we can consider what happens when a 4 hour cycle (as in *E*_2_) is split into asymmetric periods of RH and re-oxygenation by setting 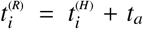 and 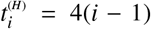 in Eq. (21), where *t_a_* ∈ (0, 4). If *t_a_* > 2, then the cells spend more time exposed to low oxygen levels (*c* < *c*\*) than to normal values (*c* > *c*\*); the opposite holds when 0 < *t_a_* < 2. We run numerical simulations up to *t* = 28 hr and report predictions of the true cell fractions, *f_m_* with *m* ∈ {1, *s*, 2} and total cell number, *N*, at 5 time points.

The results from these numerical simulations are summarised in Fig. 22. We observe that, at all time points considered, the larger the value of *t_a_* the lower is the total number of cells, *N*, (see Fig. 22(d)). When *t_a_* = 2.5*hr* (see light green dots), DNA synthesis is so slow that at the final t ime ( *t* = 28 hr) only a small fraction of cells has completed duplication and entered the *G*2/*M* phase. As a result, the fraction *f_s_* is larger than for the smaller values of *t_a_* while the fraction *f*_2_ is smaller. In this case, the variability in the model predictions remains small (even smaller than for the case *t_a_* = 2 hr which was used for the *in vitro* experiments). Therefore, our model predicts that, on short time scales (≈ 30 hr), cycles with longer periods of RH than re-oxygenation favour the accumulation of cells in the S phase. In particular, exposing cells to cycles with *t_a_* = 2.5 for 20 hours may be sufficient to synchronise them in the S phase of the cell-cycle. By contrast, when we decrease the value of *t_a_*, our model predicts only a 5% increase in the fraction of cells in the S phase. In line with our intuition, as *t_a_* decreases, fewer cells are impacted by cyclic levels of oxygen and, therefore, their distribution is more concentrated at specific phases of the cell-cycle. We note, however, that for smaller values of *t_a_* (e.g., *t_a_* = 1 or *t_a_* = 1.25), there is greater uncertainty in the predictions of *f*_1_ and *f*_2_ at *t* = 28 hr, due to the uncertainty in whether the *C*_2_ checkpoint will be activated or not (results not shown). This, again, hints at the need to refine our parameter estimates by performing new experiments (such as the cycle experiment in Fig. 20) to obtain more accurate predictions for environmental conditions that deviate from those used here to calibrate our model.

**Figure 22:**
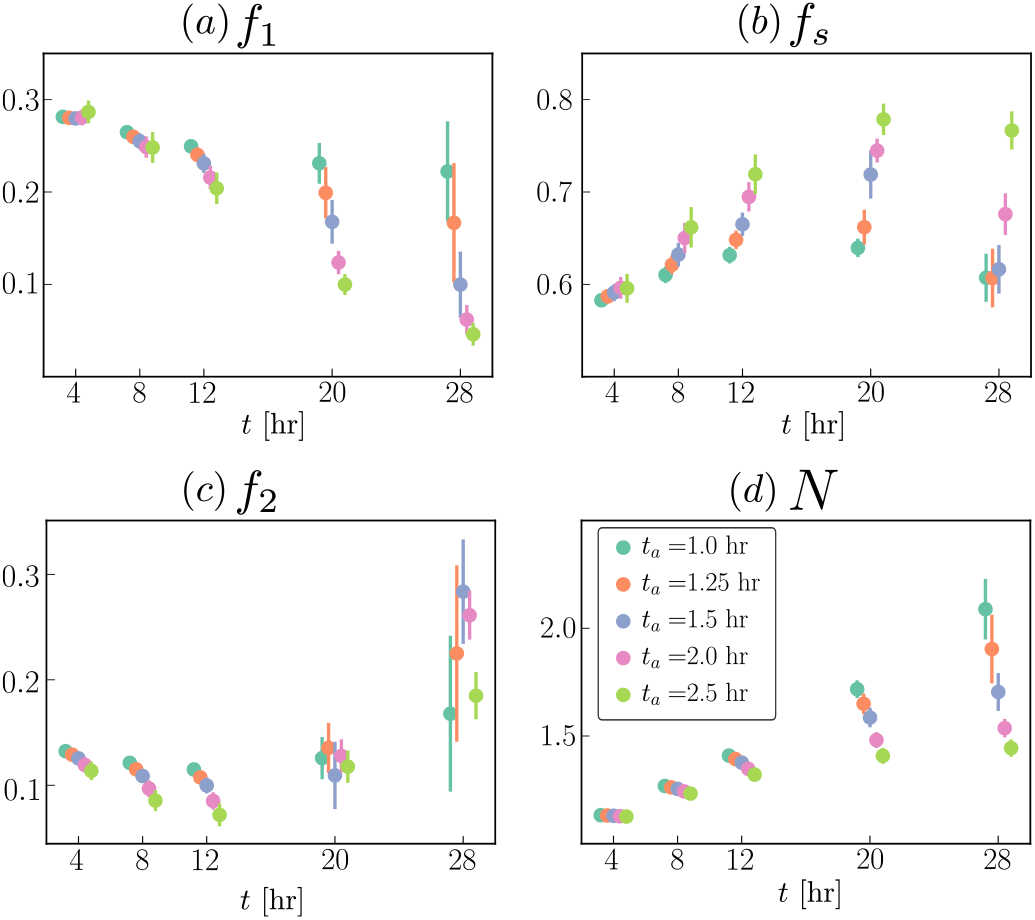
Numerically-estimated values of the cell fractions *f_m_* and cell number *N* for different cyclic protocols. We report values of the variables at 5 time points; for each variable we report the expected value based on the estimated posterior (coloured dots) and the corresponding 68%-confidence interval indicated by the vertical lines. Different colours correspond to different oxygen dynamics, *i.e.*, a different choice of *t_a_*.

## 8. Discussion and Future Work

In this paper, we have presented a five-compartment model for the cell-cycle which accounts for cell response to variable oxygen levels. We have focussed on the impact of dNTP shortages (in conditions of radio-biological hypoxia) on rates of DNA synthesis. This was achieved by introducing a variable rate, *v*(*t*), of DNA synthesis and allowing transient cell-cycle arrest in the late G1 phase by transition to a checkpoint compartment *C*_1_. A second checkpoint compartment, *C*_2_, accounts for cell-cycle arrest in the G2 phase due to accumulation of replication stress and damage. Under constant oxygen-rich conditions the model reduces to a linear, three-compartment model and analytic expressions for the long time dynamics can be derived (see §4). This analysis predicts that, in the absence of competition, cells evolve to a regime of balanced exponential growth, a result which is consistent with other cell-cycle models [8, 10, 13, 21, 54]. The main novelty of the work is the investigation of the cell-cycle dynamics in cyclic hypoxia (see §5). We show first that the model can recapitulate the experimental data from [5]. We then explore different oxygen dynamics and, in so doing, show different ways in which cyclic hypoxia can dis-regulate cell-cycle dynamics, and lead to a redistribution of cells across the phases of the cell cycle. Further, we identify scenarios in which cyclic hypoxia leads to complete inhibition of proliferation and scenarios in which proliferation is only slowed down. This is of relevance when thinking about cell-cycle specific treatment, for which changes in cell-cycle distribution (even if they are transient), can have a large impact on treatment efficacy. However, in order to use our model as a predictive tool, accurate and robust predictions are needed. In the remainder of the paper, we therefore showed how our modelling framework can be used to predict cell-cycle dynamics and inform the design of *in vitro* experiments (see §3). We started by deriving a class of candidate models 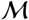 based on Eqs. (17) by systematically decreasing model complexity (i.e., the number of unknown parameters). Here our aim was to test different assumptions on the mechanisms driving cell response to cyclic hypoxia. In §6, we used Bayesian model selection to identify the best candidate model from 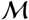 and showed that this model can indeed recapitulate the dynamic data from [5]. Furthermore, by constructing our class of models and applying Bayesian modelling selection we were able to systematically show that the inclusion of the *C*_1_ and *C*_2_ compartments in our model is necessary for it to capture the cell-cycle dynamics observed experimentally. In §7, we used our calibrated model to revisit the results from §5 where we account for uncertainty in parameter estimates. While the model makes precise predictions on short time scales (*t* ≈ 30 hr), we observed a large uncertainty in the cell-cycle dynamics at longer times. We therefore discussed how our model could be used to inform the efficient and effective design of future experiments to refine our parameter estimates, as well as to validate predictions of the calibrated model.

In this work we showed that our model can recapitulate the response to cyclic hypoxia of a specific cancer cell line (i.e., RKO cancer cell line). For this cell line, our model predicts that both constant and cyclic radio-biological hypoxia (RH) perturb the cell-cycle dynamics, but in different ways. In constant RH, cells tend to accumulate in the late G1 phase and proliferation is rapidly alted. During cyclic RH, we predict instead a more diverse range of responses depending on the oxygen dynamics. We can identify regimes where proliferation is inhibited due to accumulation of cells in the G2/M phase. By contrast, when the duration of the re-oxygenation phase is increased, population growth is only mildly slowed down and the fractions of cells in S phase is up-regulated. It would be interesting to test whether our findings can be generalised to other cell lines and how much variability there is in their response to similar cyclic protocols. Based on these observations, an interesting future research direction emerges, namely, investigating the role that cyclic RH can have on the response to cell-cycle dependent treatment, such as radiotherapy. Given that cyclic RH can change the distribution of cells around the phases of the cell-cycle, we expect a differential response to radiotherapy. This could be investigated by extending our model to include radiotherapy and to account for changes in radio-sensitivity in different phases of the cell-cycle. From this point of view, a natural question is whether cell-cycle redistribution is sufficient to explain the increase in radio-resistance due to cyclic hypoxia as reported in the literature [35, 36, 37].

In this paper, our focus was on constructing a minimal model to describe the influence of cyclic hypoxia on the cell-cycle which could be validated against existing experimental data. This guided our assumption that the rates 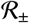 at which cells adjust their rates of DNA synthesis, *v*, to be constant. In practice, these rates may also depend on the level of damage and replication stress accumulated by the cell during cyclic hypoxia. Our model could easily be extended to account for these effects but at the cost of increasing the number of unknown model parameters. *In silico* hypothesis testing (using a Bayesian framework, as in §6), could be used to compare different modelling assumptions (i.e., constant vs variable rates), and to investigate the design of future experiments that can distinguish between the alternative mathematical models.

In several instances we have mentioned that DNA damage plays a key role in mediating cell-cycle progression and cellcycle arrest in cyclic hypoxia. However, for simplicity, we have accounted for it only implicitly. In principle, our model could be extended by introducing DNA damage as an additional structural variable to describe the cell state. While being more realistic (and of interest from a mathematical perspective), the increased model complexity would make it difficult to fit the model to experimental data. Moreover, introducing DNA damage into the model would enable us to account for radiotherapy in a more realistic manner. Analogously to re-oxygenation, radiotherapy also causes DNA damage. Such a model extension could be used to investigate whether cyclic hypoxia selects for radio-resistant clones, which are less sensitive to damage accumulation. Further model extensions could account for spatial heterogeneity, and bring the model closer to *in vivo* conditions. In this light, we view our work as a first step towards developing a theoretical framework for investigating cyclic hypoxia and its effect on cell-cycle progression, particularly in the context of solid tumours.

## Appendix A. Exponential steady state uniqueness

Starting from Eqs. (19), we note that the evolution of the number of cells, *N*, decouples from the equations for *G*_1_ and *G*_2_ so that we can consider the reduced model:

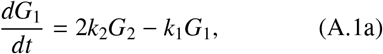

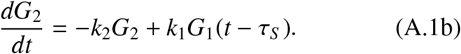

Since Eqs. (A.1) form a system of linear delay differential equations, their solutions can be written as a superposition of exponential functions *e*^Λ*t*^ with corresponding eigenvalue 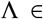 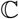. We therefore know that the long time behaviour of the system will be dominated by the eigenvalue Λ with largest real part, here denoted by λ. Unlike for ordinary differential equations, the number of eigenvalues Λ for a delay differential equation is infinite and they are defined by the characteristic equation:

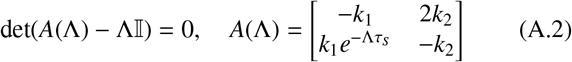

where 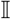 is the identity matrix in ℝ^2×2^. Evaluating the determinant explicitly we obtain the following transcendental equation, whose roots correspond to the eigenvalues Λ of Eqs. (A.1):

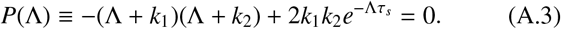

It can be shown that this system has at least one root with Re(Λ) > 0 so that the solution will blow up in time, i.e. 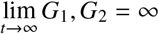. We also have that the spectrum of eigenvalues is bounded above, in the sense that there exists an upper limit to the values of Re(Λ). This is analogous to the system investigated by Crivelli et al. in [21]. Lemma 1 summarises some of their results as adapted to our model.

### Lemma 1.

For any value of k_1_ > 0, k_2_ > 0 and τ_s_ > 0, the right-most root of P(Λ) as defined by Eq. (A.3) is real and positive.

### Proof.

Let us first consider the existence of a real and positive root Λ = Λ_0_. This is straightforward to prove since *P*(Λ) is continuous and *P*(0) > 0 while lim_Λ→∞_ *P*(Λ) < 0. Given that for Λ > 0, *dP*/*d*Λ < 0, i.e. *P* is strictly monotonically decreasing, we have that the zero Λ = Λ_0_ is unique. This implies that for any choice of parameters, the trivial steady state (0, 0) is unstable.

Let us now consider the complex solution Λ = Λ_*R*_ + *i*Λ_*I*_ for the function *P* where Λ_*I*_ ∈ ℝ \ {0}:

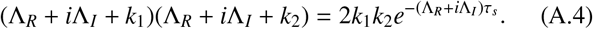

Taking absolute values of the above we obtain that:

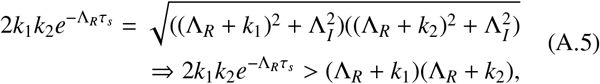

which implies that *P*(Λ_*R*_) > 0 = *P*(Λ_0_), where Λ_0_ is the unique real root of *P*(Λ). Since *P* is strictly monotonically decreasing we therefore have that Λ_*R*_ > Λ_0_. We therefore have that the rightmost eigenvalue is real and it is positive.

Based on Lemma 1, we know that for any choice of model parameters, under unperturbed growth, cells will eventually reach a regime in which they grow exponentially. This is a common result of many cell-cycle models for *in vitro* systems and it is usually referred to as balanced or asynchronous exponential growth:

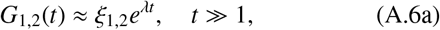

where *ξ*_1,2_ are positive constants and the character 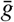 is used to indicate the asymptotic solution. Having characterised the long time behaviour of *G*_1_(*t*) and *G*_2_(*t*), let us discuss what this implies for the other model variables, i.e. the distribution *S* (*x*, *t*) and the total number of cells, *N*. Using Eq. (9) we find that, in the case of unperturbed growth and assuming *t* > τ_*s*_, the long time distribution *S* (*x*, *t*) takes the form:

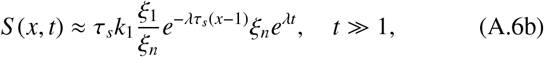

which can be written by separating the DNA and time components, as 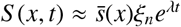. This implies that the population *P*_*s*_ also grows exponentially, 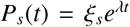 where 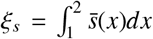. Combining this with Eq. (A.6a), we can compute the total number of cells and the cell fractions:

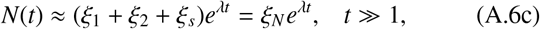

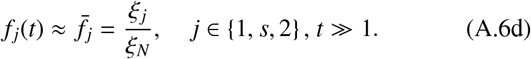

We therefore find that the long time behaviour is characterised by a stationary DNA-distribution, 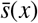 of cells in the S phase, and constant cell fractions, 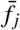. Following [54] we will denote this specific regime as the *steady phase solution* to highlight the fact that the fraction of cells in each phase of the cycle remains constant.

We now discuss how this *steady phase solution* of the model can be used to estimate the model parameters. Assume that the cells are left growing in an unperturbed environment for sufficiently long time so as to reach the regime of balanced exponential growth. Provided that we know the fraction of cells, 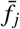, and proliferation rate, λ, of the population of cells (which can be approximated using the doubling time as given in the main text, *T_doub_* = λ^−1^ ln 2), we can uniquely identify the parameters *k*_1_, *k*_2_, τ_*s*_ in the model. Let us substitute the solution (A.6) into (19) and, upon re-writing 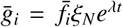, we obtain the following algebraic system:

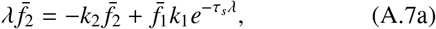

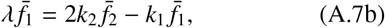

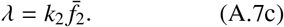

Looking at Eq. (A.7c) it is apparent that *k*_2_ is uniquely identified and it is positive. Substituting *k*_2_ into Eq. (A.7b), we obtain an equation for *k*_1_:

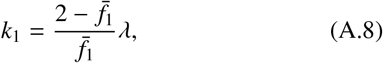

where we note that *k*_1_ > 0 since, by definition, 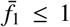. Sub-stituting now the forms of *k*_1_ and *k*_2_ into (A.7a) we obtain an expression for τ_*s*_:

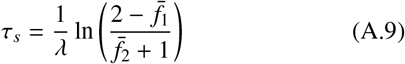

where the physical constraint 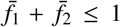 guarantees that the argument of the logarithm is always positive. Hence, given the measurements of 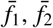, τ_*s*_ we can uniquely identify the constant parameters appearing in the model (19) for unperturbed growth.

## Appendix B. Experimental data

Here we present the raw-data for the cell-cycle dynamics in RKO cancer cells as measured in the experiments from [5] discussed in §2 of the main text. Table B.6 summarises the value of the cell fractions in normoxia (*E*_0_), while Table B.7 and Table B.8 represent the time series data obtained when cells are exposed to constant (*E*_1_) and cyclic (*E*_2_) RH, respectively.

**Table B.6:** Raw experimental data for the stationary cell-cycle dynamics of RKO cancer cells when exposed to normoxia (*E*_0_) for sufficiently long time.

| $F_1$ | $F_s$ | $F_2$ |
| --- | --- | --- |
| 0.291222313 | 0.557013946 | 0.151763741 |
| 0.297764120 | 0.533750132 | 0.168485748 |
| 0.276288660 | 0.602061856 | 0.121649485 |
| 0.391933816 | 0.475698035 | 0.132368149 |
| 0.270000000 | 0.540000000 | 0.190000000 |
| 0.353889645 | 0.498481834 | 0.147628521 |
| 0.296387429 | 0.515714126 | 0.187898445 |
| 0.289529012 | 0.571414043 | 0.139056945 |
| 0.291841004 | 0.560669456 | 0.147489540 |

**Table B.7:** Raw experimental data for the cell-cycle dynamics of RKO cancer cells when exposed to constant RH (*E*_1_).

| Time [hr] | $F_1$ | $F_s$ | $F_2$ |
| --- | --- | --- | --- |
| 4 | 0.491356731 | 0.297537978 | 0.207438449 |
| 4 | 0.497400877 | 0.36815819 | 0.131485068 |
| 8 | 0.513138837 | 0.334160977 | 0.147941237 |
| 8 | 0.563423645 | 0.325328407 | 0.104679803 |

**Table B.8:** Raw experimental data for the cell-cycle dynamics of RKO cancer cells when exposed to cyclic RH (*E*_2_).

| Time [hr] | $F_1$ | $F_s$ | $F_2$ |
| --- | --- | --- | --- |
| 4 | 0.330630725 | 0.546067179 | 0.120037907 |
| 4 | 0.266513378 | 0.632211538 | 0.098035117 |
| 4 | 0.297175141 | 0.534011299 | 0.164971751 |
| 8 | 0.220709571 | 0.731332508 | 0.044657591 |
| 8 | 0.191676374 | 0.754315366 | 0.050937202 |
| 20 | 0.159383033 | 0.702313625 | 0.133676093 |
| 20 | 0.093743547 | 0.777617179 | 0.125954987 |
| 20 | 0.144138373 | 0.711189408 | 0.140935298 |
| 20 | 0.100658314 | 0.726587386 | 0.168825653 |
| 20 | 0.184611856 | 0.635248251 | 0.171998624 |
| 20 | 0.145497026 | 0.713785047 | 0.13700085 |
| 28 | 0.060079664 | 0.712547024 | 0.21132994 |
| 28 | 0.044509303 | 0.647087438 | 0.29916089 |
| 28 | 0.03960499 | 0.676611227 | 0.280665281 |

## Appendix C. MCMC sampling algorithm and results

In order to estimate the posterior distribution (see Eq. (24a)), we use Markov Chain Monte Carlo (MCMC) methods, employing the freely available implementation in the python package PINTS [19]. As suggested by Johnstone et al. [38], prior to starting our MCMC routine, we compute a good initial guess by maximizing the likelihood function 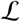 (see Eq. (23b)). Based on Eq. (24a) and the choice of a uniform prior, maximising the posterior is equivalent to maximising the likelihood function. Sampling random initial guesses from π_*pr*_, we solve the optimisation problem for the log-likelihood using the `CMA-ES` algorithm [19, 34]. We then use the output of the optimization routine to initiate the MCMC simulation (we compute, in total, three chains). We sample from the posterior distribution by using `HaarioBardenet MCMC`, which is a Metropolis-Hastings algorithm with adaptive covariance, where the first 8000 iterations are performed without adaptation (as suggested by [38]). We compute up to 30000 iterations for each chain and discard the first 10000 as “*warm-up*”. As in [20], we assess the convergence of the MCMC chains by estimating 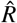 [41, chapter 13], where we accept the sampled posterior if 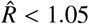 (note that 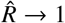 as the algorithm converges).

In Figs. C.23–C.26, we show the estimated posterior distributions for our class of models *M* (see Table 3). Summary statistics of the marginal posterior distributions are listed in Table C.9. For the point estimates of parameter values in §5, we use the mean of the marginal posterior distributions (see Table C.9).

**Table C.9:** Summary statistics for the marginal posterior distributions for the family of models *M* listed in Table 3 in §5.2. We here report the mean and standard deviation together with the quantiles (*Q_i_*) of the distribution. The last column shows the value of 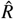 used to estimate the convergence of the MCMC algorithm, where convergence corresponds to 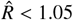.

| Model | | mean | std | $Q_{25\%}$ | $Q_{50\%}$ | $Q_{75\%}$ | $\hat{R}$ |
| --- | --- | --- | --- | --- | --- | --- | --- |
| $\mathcal{M}_0$ | $\alpha_1$ | 1.57 | 0.50 | 1.17 | 1.55 | 1.96 | 1.01 |
| | $\mathcal{R}_+$ | 0.25 | 0.11 | 0.17 | 0.24 | 0.32 | 1.01 |
| | $\mathcal{R}_-$ | 0.28 | 0.10 | 0.21 | 0.27 | 0.34 | 1.01 |
| | $p$ | 0.83 | 0.11 | 0.77 | 0.85 | 0.91 | 1.02 |
| | $q$ | 0.89 | 0.06 | 0.86 | 0.90 | 0.94 | 1.01 |
| | $T$ | 16.89 | 1.16 | 16.21 | 17.02 | 17.74 | 1.01 |
| | $K_2$ | 1.04 | 0.55 | 0.59 | 1.06 | 1.51 | 1.00 |
| | $\sigma_1$ | 0.04 | 0.01 | 0.03 | 0.04 | 0.04 | 1.01 |
| | $\sigma_2$ | 0.05 | 0.01 | 0.03 | 0.05 | 0.05 | 1.01 |
| $\mathcal{M}_1$ | $\alpha_1$ | 1.53 | 0.50 | 1.13 | 1.50 | 1.94 | 1.00 |
| | $\mathcal{R}_+$ | 0.26 | 0.12 | 0.17 | 0.25 | 0.33 | 1.00 |
| | $\mathcal{R}_-$ | 0.30 | 0.13 | 0.22 | 0.28 | 0.58 | 1.00 |
| | $p$ | 0.83 | 0.11 | 0.76 | 0.85 | 0.91 | 1.00 |
| | $q$ | 0.90 | 0.06 | 0.86 | 0.91 | 0.95 | 1.00 |
| | $T$ | 16.93 | 1.18 | 16.21 | 17.05 | 17.78 | 1.00 |
| | $\sigma_1$ | 0.04 | 0.01 | 0.03 | 0.04 | 0.04 | 1.00 |
| | $\sigma_2$ | 0.05 | 0.01 | 0.04 | 0.05 | 0.05 | 1.00 |
| $\mathcal{M}_2$ | $\mathcal{R}_+$ | 0.11 | 0.08 | 0.05 | 0.10 | 0.15 | 1.01 |
| | $\mathcal{R}_-$ | 0.14 | 0.06 | 0.10 | 0.13 | 0.17 | 1.00 |
| | $p$ | 0.71 | 0.16 | 0.59 | 0.71 | 0.83 | 1.01 |
| | $T$ | 15.29 | 0.87 | 14.75 | 15.27 | 15.86 | 1.01 |
| | $\sigma_1$ | 0.14 | 0.02 | 0.14 | 0.15 | 0.19 | 1.00 |
| | $\sigma_2$ | 0.04 | 0.01 | 0.04 | 0.04 | 0.05 | 1.00 |
| $\mathcal{M}_3$ | $\alpha_1$ | 1.63 | 0.53 | 1.22 | 1.66 | 2.08 | 1.01 |
| | $\mathcal{R}_+$ | 0.21 | 0.19 | 0.08 | 0.16 | 0.27 | 1.04 |
| | $\mathcal{R}_-$ | 0.53 | 0.31 | 0.32 | 0.45 | 0.66 | 1.03 |
| | $q$ | 0.93 | 0.05 | 0.89 | 0.93 | 0.97 | 1.01 |
| | $\sigma_1$ | 0.04 | 0.01 | 0.04 | 0.04 | 0.05 | 1.01 |
| | $\sigma_2$ | 0.12 | 0.02 | 0.11 | 0.12 | 0.14 | 1.01 |

**Figure C.23:**
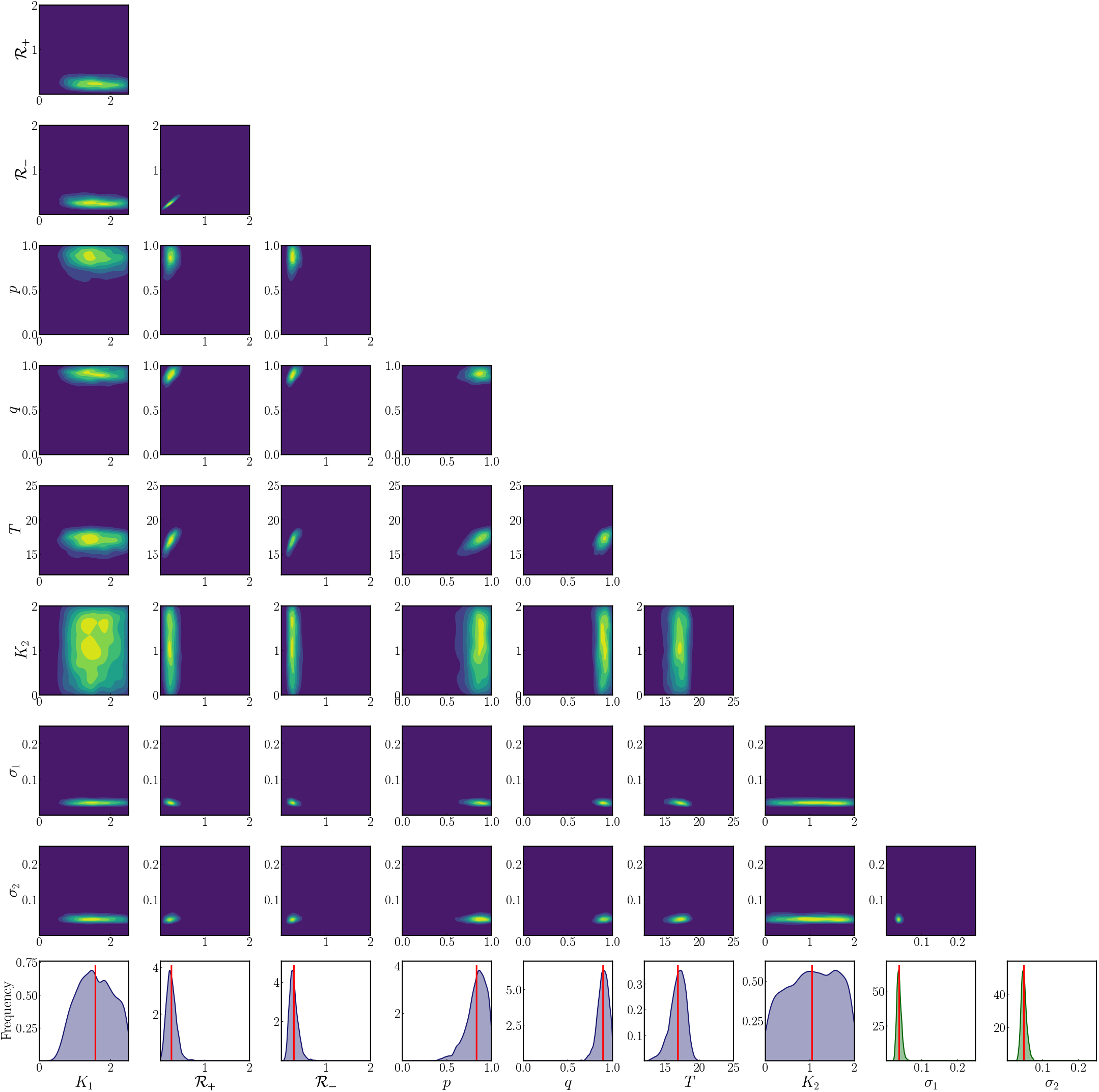
Approximation of the joint (surface plot) and marginal (last row) posterior distributions for model 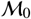 from Table 3. The distributions were obtained by using the MCMC samples generated as discussed in the text. In the surface plot for joint distributions, yellow areas correspond to higher posterior probability in contrast to the blue areas which correspond to low probability. In the last row, the red vertical line indicates the mean of the marginal posterior distribution as reported in Table C.9, where additional summary statistics extrapolated from the marginal distribution are also given.

**Figure C.24:**
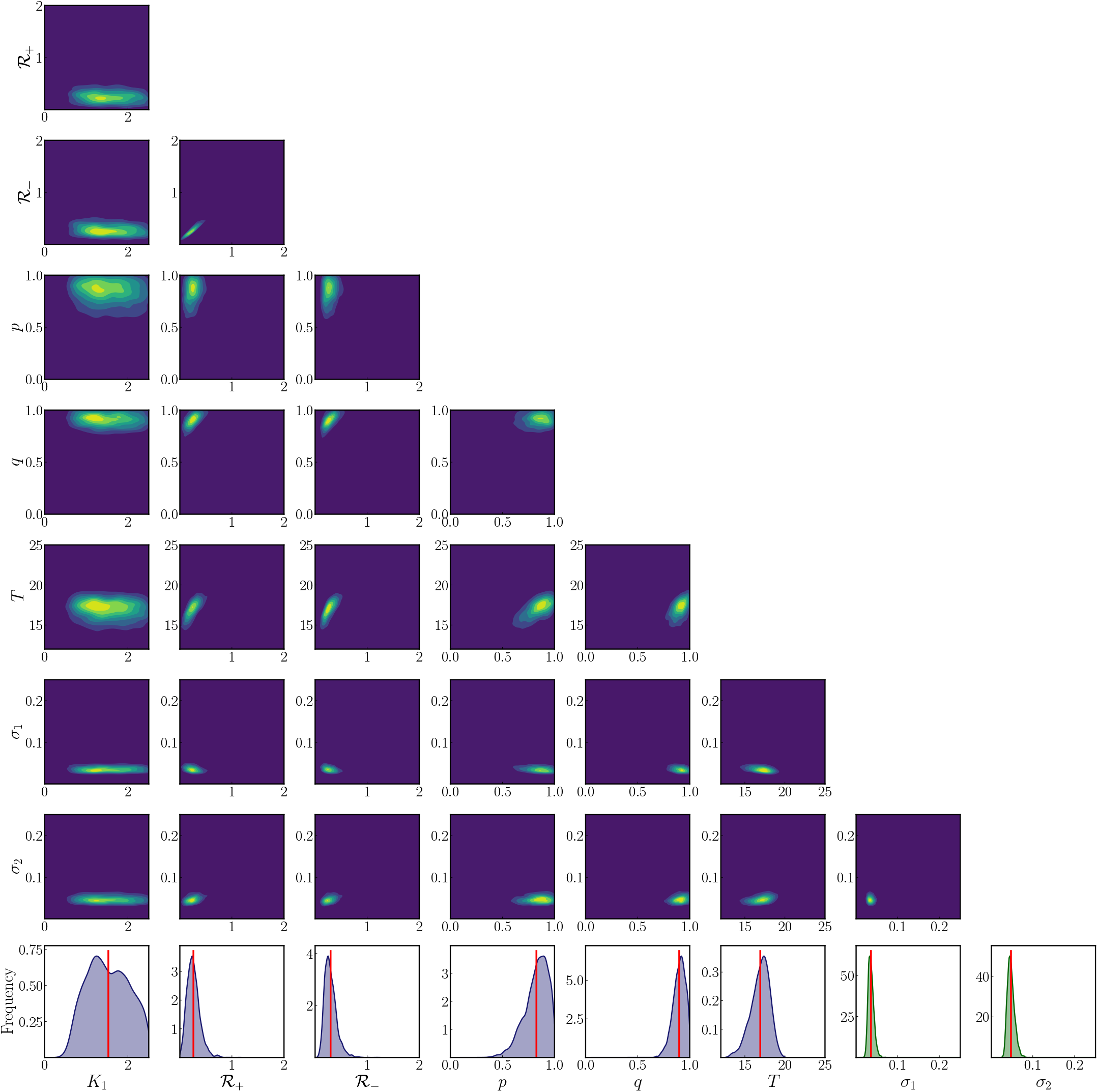
Approximation of the joint (surface plot) and marginal (last row) posterior distributions for model 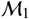 from Table 3. The distributions were obtained by using the MCMC samples generated as discussed in the text. In the surface plot for joint distributions, yellow areas correspond to higher posterior probability in contrast to the blue areas which correspond to low probability. In the last row, the red vertical line indicates the mean of the marginal posterior distribution as reported in Table C.9, where additional summary statistics extrapolated from the marginal distribution are also given.

**Figure C.25:**
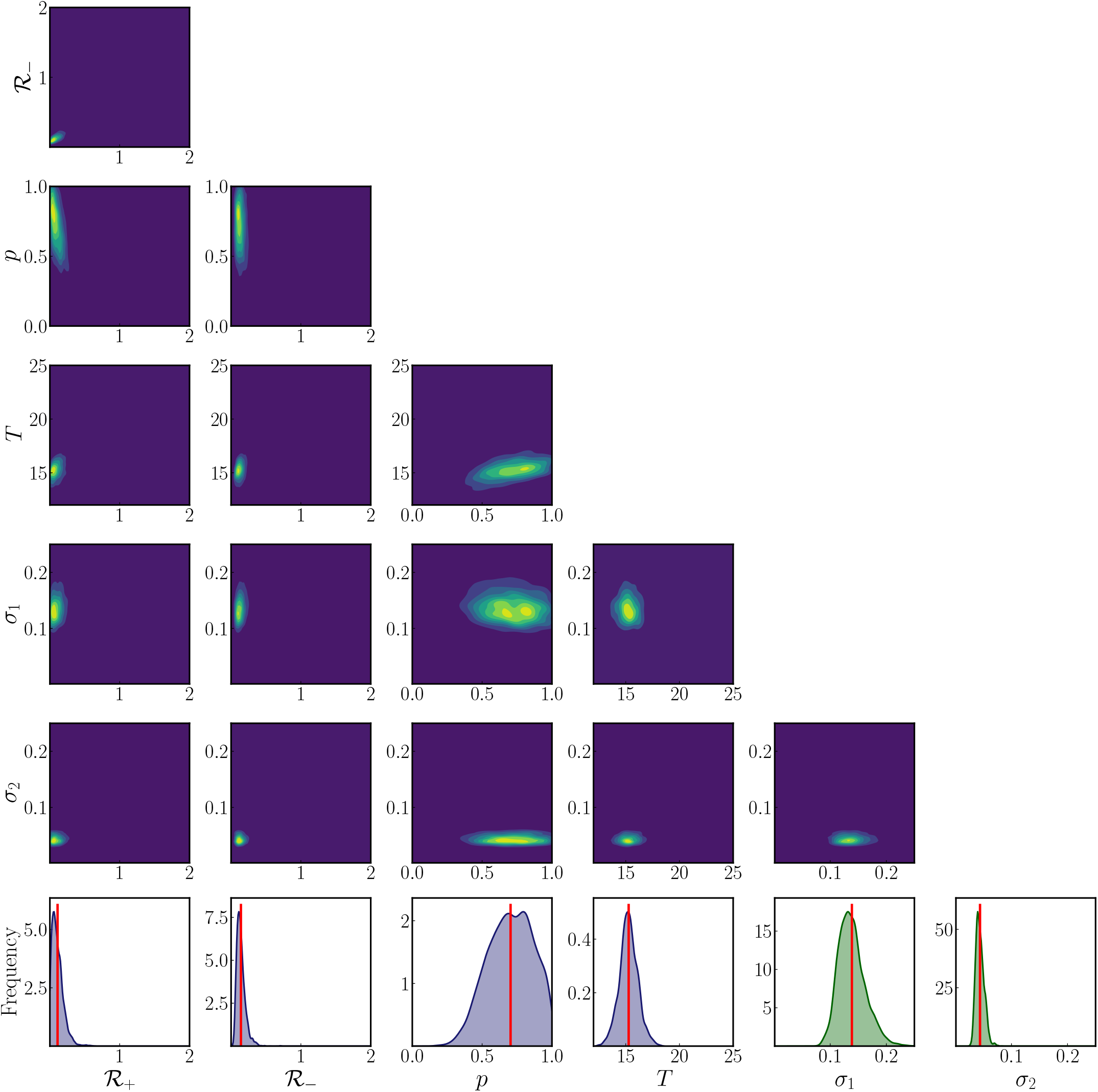
Approximation of the joint (surface plot) and marginal (last row) posterior distributions for model 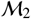 from Table 3. The distributions were obtained by using the MCMC samples generated as discussed in the text. In the surface plot for joint distributions, yellow areas correspond to higher posterior probability in contrast to the blue areas which correspond to low probability. In the last row, the red vertical line indicates the mean of the marginal posterior distribution as reported in Table C.9, where additional summary statistics extrapolated from the marginal distribution are also given.

**Figure C.26:**
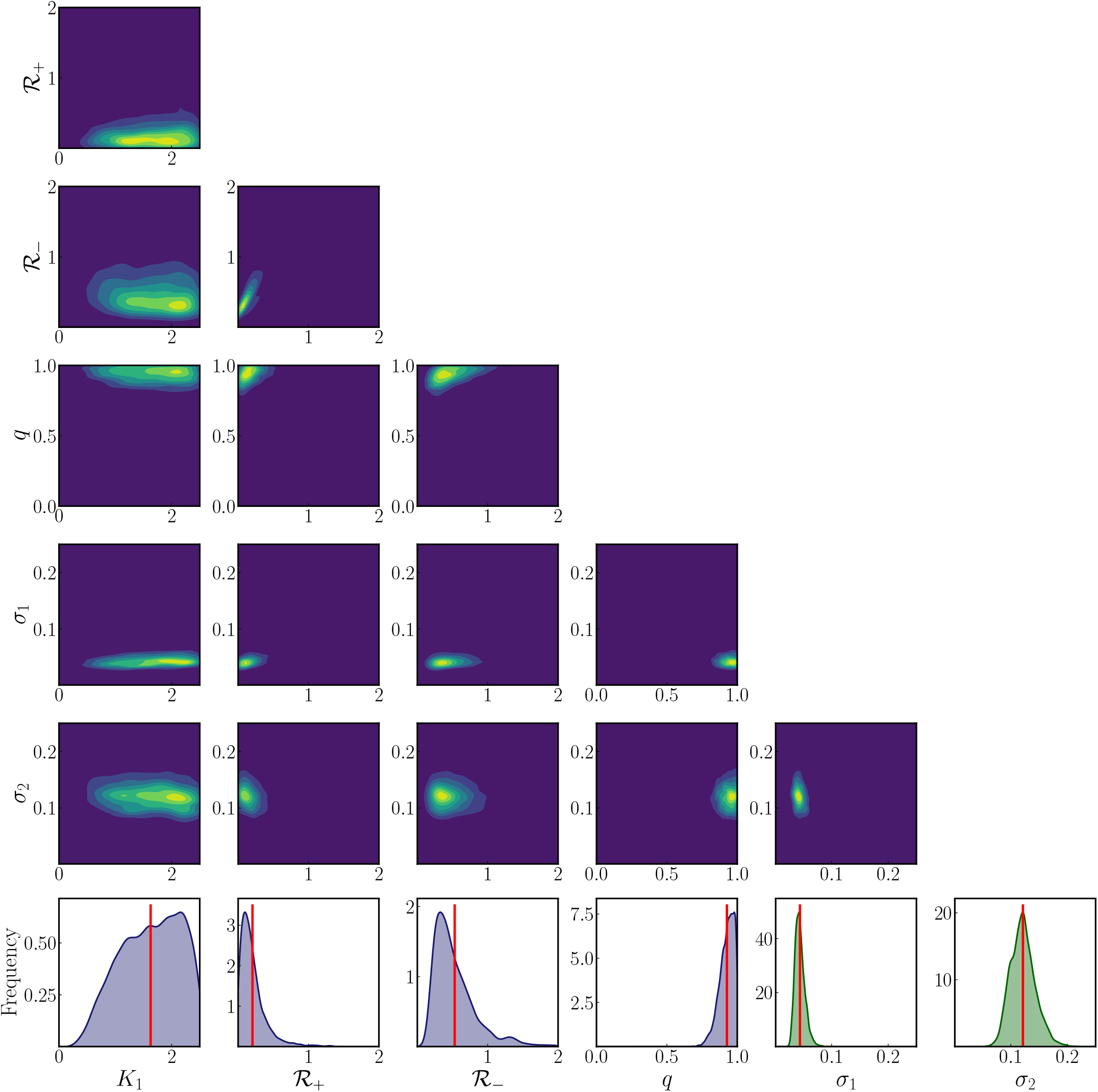
Approximation of the joint (surface plot) and marginal (last row) posterior distributions for model 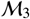 from Table 3. The distributions were obtained by using the MCMC samples generated as discussed in the text. In the surface plot for joint distributions, yellow areas correspond to higher posterior probability in contrast to the blue areas which correspond to low probability. In the last row, the red vertical line indicates the mean of the marginal posterior distribution as reported in Table C.9, where additional summary statistics extrapolated from the marginal distribution are also given.

